# Axial spondyloarthritis patients have altered mucosal IgA response to oral and fecal microbiota

**DOI:** 10.1101/2021.07.15.452558

**Authors:** Tejpal Gill, Patrick Stauffer, Mark Asquith, Ted Laderas, Tammy M. Martin, Sean Davin, Matthew Schleisman, Claire Ramirez, Kimberly Ogle, Ingrid Lindquist, Justine Nguyen, Stephen R. Planck, Carley Shaut, Sarah Diamond, James T. Rosenbaum, Lisa Karstens

## Abstract

**Objective:** To investigate whether axial spondyloarthritis (AxSpA) patients have an altered immunoglobulin A (IgA) response in the gut and oral microbial communities.

**Methods:** We performed 16S rRNA gene (16S) sequencing on IgA positive (IgA^+^) and IgA negative (IgA^−^) fractions (IgA-SEQ) from feces (n=17 AxSpA; n=14 healthy) and saliva (n=17 AxSpA; n=12 healthy), as well as on IgA-unsorted fecal and salivary samples. PICRUSt2 was used to predict microbial metabolic potential in AxSpA patients and healthy controls (HCs).

**Results:** IgA-SEQ revealed enrichment of several microbes in the fecal (*Akkermansia, Ruminococcaceae, Lachnospira*) and salivary (*Prevotellaceae, Actinobacillus*) microbiome in AxSpA patients as compared with HCs. Fecal microbiome from AxSpA patients showed a trend towards increased alpha diversity of the IgA^+^ fraction and decreased diversity in the IgA^−^ fraction in comparison with HCs, while the salivary microbiome exhibits a significant decrease in alpha diversity in both IgA^+^ and IgA^−^ fractions. Increased IgA coating of *Clostridiales Family XIII* correlated with disease severity. Inferred metagenomic analysis suggests perturbation of metabolites and metabolic pathways for inflammation (oxidative phosphorylation, glutathione metabolism) and metabolism (propanoate and butanoate metabolism) in AxSpA patients.

**Conclusions:** Analyses of fecal and salivary microbes from AxSpA patients reveal distinct populations of immunoreactive microbes using novel IgA-SEQ approach, which were not captured by comparing their relative abundance with HCs. Predictive metagenomic analysis revealed perturbation of metabolites/metabolic pathways in AxSpA patients. Future studies on these immunoreactive microbes may lead to better understanding of the functional role of IgA in maintaining microbial structure and human health.

## Introduction

Axial spondyloarthritis (AxSpA) is an immune-mediated inflammatory arthritis, which affects the sacroiliac and spinal joints, and is associated with microscopic lesions or inflammation in the gut (1). It includes ankylosing spondylitis (AS), in which radiographic evidence of inflammation in spine or sacroiliac joints is observed, and non-radiographic AxSpA in which radiographic changes are not observed. While the genetic association of AS with the major histocompatibility molecule HLA-B27 has been known for almost five decades (2), the mechanistic link remains elusive. Like several other complex polygenic diseases such as inflammatory bowel disease (IBD), diabetes mellitus and multiple sclerosis, gene-environment interactions and particularly host-microbiota interactions have been centrally implicated in pathogenesis (reviewed in (3)). The latter is well supported by the observations that patients with AS have a dysbiotic gut microbiota and over half exhibit subclinical bowel inflammation (4–7).

Previously, we have shown that HLA-B27 expression perturbs the gut microbiota in an experimental model of spondyloarthritis (SpA) (8, 9). In this model, transgenic rats expressing multiple copies of human HLA-B27 in conjunction with human β2-microglobulin (HLA-B27 TG) develop spontaneous bowel and joint disease (10). Enhanced mucosal immune responses to the gut microbiota strongly correlate with disease severity in these animals. For instance, HLA-B27 TG rats with arthritis elicit a stronger microbiota-specific IgA response (increased frequency of IgA-coated microbes) than control HLA-B7 TG animals without joint disease (8). Recently, we have shown that healthy individuals carrying the HLA-B27 allele also have altered gut microbial composition (11). While multiple studies employing 16S rRNA (16S) and metagenomic sequencing have revealed gut microbial dysbiosis associated with various spondyloarthropathies (12–17), identification of a causal microbe has been elusive. Compositional analysis of the microbial community may not be sufficient to determine the interactions between microbes and the host immune response. To identify such bacteria that preferentially affect disease susceptibility and severity, we employed IgA-sequencing (IgA-SEQ), which couples flow cytometry sorting of IgA-coated bacteria and 16S sequencing, to identify IgA coated microbes (18). This technique has been used to determine pathologically relevant microbes in Crohn’s disease (CD)-associated SpA (19). AS and CD share several aspects of clinical overlap including bowel, eye, and joint disease (20). However, this type of investigation of the mucosal immune response has not yet been reported in AxSpA patients.

In this study, we evaluated IgA coating (IgA-SEQ) and traditional 16Ssequencing of microbes in the saliva and feces from AxSpA patients and healthy controls (HCs). IgA-SEQ analyses identified potentially immune-targeted microbes that were not observed to be differentially abundant using traditional 16S sequencing. In addition, differential IgA coating on some microbes was associated with disease activity. Using a computational approach to predict the microbial function, we found that the IgA^+^ fraction in the fecal or salivary samples showed perturbation of many predictive metabolites and metabolic pathways in AxSpA patients. These potentially immune reactive microbes may reveal novel host-microbial interactions in health and disease. Taken together, this study adds to our understanding of HLA-B27-associated host immune response and its interaction with the gut microbes in AxSpA.

## Patients and Methods

### Study design and participants

We performed a prospective cohort study at the Oregon Health & Science University (OHSU). Subjects were excluded from all cohorts if they were younger than age 18 years, pregnant, had a history of prior intestinal surgery or colon cancer, or had antibiotic use 6 months prior to sample collection. This study was approved by the Institutional Review Board at OHSU and written informed consent was obtained from all participants.

#### AxSpA cohort

Subjects with a diagnosis of AxSpA were recruited from either the OHSU rheumatology clinic or via the Spondylitis Association of America (SAA). The SAA is a patient advocacy group with physicians on their board, and members with AS are highly engaged and informed. Medical records were reviewed by a rheumatologist (JTR) to confirm a diagnosis of AS based on the modified NY Criteria (21). A total of 29 subjects with AS were enrolled (12 subjects provided fecal samples, 12 provided salivary samples and 5 subjects provided both fecal and salivary samples). Of these, it was not possible to obtain medical records from external clinics for 10 subjects, all recruited from the SAA (2 subjects provided fecal samples, 7 subjects provided salivary samples and 1 subject provided both a fecal and salivary sample). These subjects self-reported specifically that they had received a diagnosis of AS, thus, in the absence of radiographic evidence for these 10 subjects we were comfortable in grouping this cohort as AxSpA. The Bath ankylosing spondylitis disease activity index (BASDAI) was administered to subjects with AxSpA. Their current medications, including biologics, as well as clinical HLA-B27 testing, if known, was also documented.

#### Samples

All subjects enrolled at OHSU provided a blood sample for genomic DNA extraction and subsequent HLA-B typing. Subjects recruited through the SAA shipped fecal and/or salivary samples to our lab using provided next-day shipping materials. For microbiome analysis, fecal and salivary samples were snap frozen upon collection and stored at −80°C until further analyses. All processing thereafter was performed blinded to patient phenotype.

#### HLA-B typing

Genomic DNA was extracted from anti-coagulated blood by the salting out method performed in a shared core resource at the Casey Eye Institute, OHSU. The LABType XR HLA-B SSO typing kit from One Lambda Thermo Fisher (RSSOX1B) was used according to manufacturer’s instructions in the OHSU Laboratory of Immunogenetics and Transplantation.

### Microbial community analysis

To determine the microbial community structure, genomic DNA from the salivary and fecal samples was isolated using the methods described previously (22). Briefly, genomic DNA was isolated using the Power Soil DNA Isolation Kit (MoBio Laboratories Inc., Carlsbad, CA) according to manufacturer’s instructions. Amplification of the 16S ribosomal RNA genes was performed using the 515–806 primers specified by the Earth Microbiome Project (https://www.earthmicrobiome.org/). Sequencing of the V4 region was accomplished with Illumina MiSeq using barcoded primers (23). For IgA-SEQ, fecal samples were homogenized in phosphate buffered saline (PBS), supernatant was collected and washed in PBS with 1% fetal bovine serum (FBS) and incubated with blocking buffer (PBS with 20% rat serum) for 20 min. Samples were stained with either mouse anti-human IgA (Miltenyi 130-093-128) or isotype mouse IgG1 control (Miltenyi 130-092-212) and sorted on a FACS Aria (BD Biosciences). Genomic DNA from IgA^+^ and IgA^−^ fractions were isolated and the V4 region of the16S rRNA gene was sequenced as described above.

Sequencing data were demultiplexed and processed using the DADA2 package in R. This included quality filtering, denoising using the DADA2 algorithm, chimera detection and removal, and tabulation into amplicon sequence variants (ASVs). Taxonomy assignment was performed using the RDP classifier (for Phylum to Genus annotations) and exact matching (for Species level annotations) with the Silva database (v. 132). Phylogenetic trees were generated using phangorn and decipher. Further analyses were performed using microbiome data analysis packages in R (phyloseq, vegan, MicrobiotaProcess) (24). IgA-SEQ data were subjected to decontam to remove potential contaminant ASVs (25), and analyses were performed using in-house written scripts, available on Github (https://github.com/KarstensLab/igaseq). Samples that had 7,500 reads per sample in both the IgA^−^ and IgA^+^ fractions and IgA index was calculated for each genus or ASV (26) were used. Alpha diversity was evaluated by using the number of observed genera/ASVs, the Shannon Index, and Inverse Simpson Index. Beta diversity was evaluated using the phylogenetic tree-based metric, UniFrac metric (27) and visualized with Principal Coordinate Analysis (PCoA).

### Predictive Metagenomic Analysis

To infer the metagenome of oral and gut microbiota from 16S rRNA amplicon sequences obtained, we employed PICRUSt2 (Phylogenetic Investigation of Communities by Reconstruction of Unobserved States 2)(28). PICRUSt2 estimates the metagenomic contribution of gene families in bacteria and archaea using 16S rRNA marker gene data, while taking into account the copy number of 16S rRNA gene. Functional potential was estimated by mapping 16S rRNA sequence-based data to predicted functional annotation via alignments and hidden state prediction models (31), using default parameters. We used the quantitative abundance counts that mapped the Kyoto Encyclopedia of Genes and Genomes (KEGG) ortholog (KO) functional categories and focused on KEGG hierarchical KEGG level 3 (pathways) and individual KOs. Overlap between KEGG predictive genes is determined between various groups, and area proportional Euler plots were created using Eulerr package in R environment.

### Statistical analysis

All statistical analyses were performed in R environment. For 16S sequencing and IgA-SEQ analyses, non-parametric tests (Wilcoxon Rank Sum) were used to assess within and between group differences for the alpha diversity measures. Differences in overall microbial community structure were evaluated using Permutational Multivariate Analysis of Variance (PERMANOVA). LEfSe was used to determine the presence of significant discriminant taxa by pairwise comparison (29). For PICRUSt2 analyses, LEfSe was used for significance testing of the abundance counts of KEGG level 3 and KOs (29). LEfSe was performed with class representing diagnosis (AxSpA or HC) and subclass representing patient sex, with an alpha <0.05 for the ANOVA comparing class level and with an alpha <0.1 for the Wilcoxon test at the subclass level. Comparisons were made between IgA^+^ and IgA^−^ fractions from salivary and fecal microbiome data from AxSpA patients and HCs. Multiple comparisons were adjusted for, where appropriate, using the Benjamini-Hochberg method (30).

## Results

### Clinical characteristics defining AxSpA patients

To analyze microbial differences between AxSpA subjects and HCs, we performed 16S sequencing on the unsorted microbial community and IgA-SEQ on IgA^+^ and IgA^−^ sorted fractions from the fecal and salivary samples. The fecal cohort had 31 subjects (17 AxSpA patients and 14 HCs), and the salivary cohort had 29 subjects (17 AxSpA patients and 12 HCs). While there were no significant differences in age or gender (*P*> 0.2 and 0.6 respectively) between the AxSpA and HC groups for the fecal samples, the HCs were younger (*P*= 0.017), and with less female representation (*P*= 0.014) than the AxSpA patients in the salivary cohort. However, both AxSpA patients and HCs had similar BMI in the fecal (*P*> 0.5) and salivary (*P*> 0.6) cohort. AxSpA patients had a mean disease activity (BASDAI) of 3.5 in the fecal cohort, and 2.2 in the salivary cohort. In both cohorts, 50% of these patients were on biologics such as adalimumab, etanercept or secukinumab (Table 1).

**Table 1.**
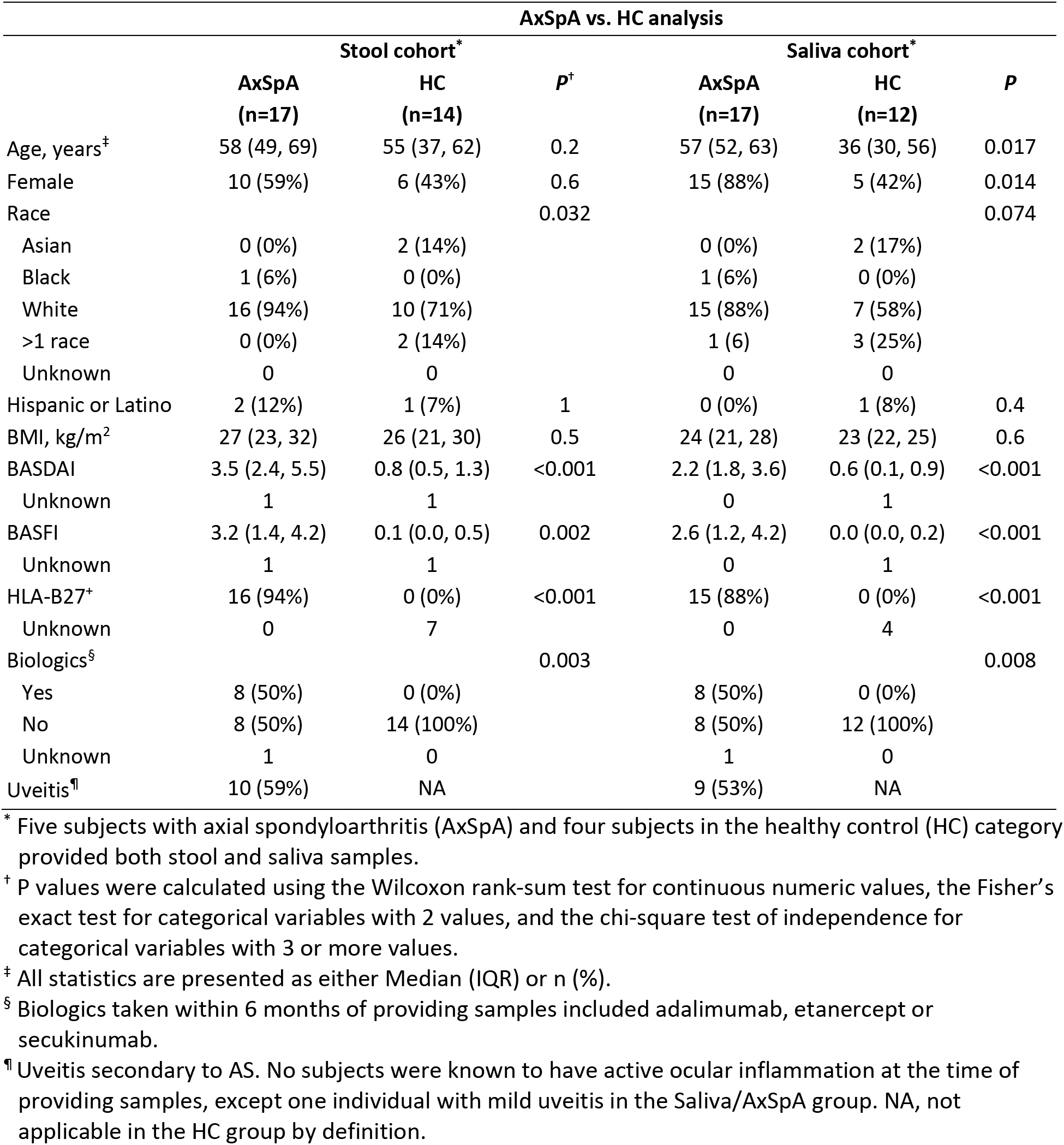
General characteristics of the subjects providing stool and saliva samples for IgA-Seq analyses.

### AxSpA patients have altered microbial composition in saliva

To characterize microbial diversity in the fecal and salivary samples from patients with AxSpA in comparison with the HCs, we performed 16S sequencing on both fecal (Figures 1A-C) and salivary (Figures 1D-F) samples. While AxSpA patients had decreased fecal microbial diversity in comparison to HCs, the differences in overall alpha and beta diversity were not significant (Figure 1A). In contrast, salivary samples from AxSpA patients showed a significant decrease in all (observed, Shannon and Inverse Simpson) measures of alpha diversity (Figure 1D) as well as differences in microbial composition (beta diversity, Figure 1E) at the genus level. We also compared individual microbial differences between AxSpA patients and HCs at the genus level for fecal and salivary samples. There were significant differences in the relative abundance of individual microbes in the fecal (e.g., *Anaerofilum, Fecalitalea, Fournierella* (Figure 1C) as well as in the salivary samples (e.g., *Megaspora, Aggregatibacter, Ruminococcaceae_UCG-014, Campylobacter, Treponema, Atopobium, Corynebacterium* and *Dialister* (Figure 1F). However, these differences no longer reached statistical significance after multiple testing correction. Similar results were seen in both fecal and salivary samples at the species level as measured using ASVs (Supplementary Figure 1A-F). Notably, there were significant difference between the salivary alpha diversity (Shannon’s Index, *P*= 0.0006) and microbial composition (*P*= 0.025) (Supplementary Figures 1D and E), but these differences were not observed in the fecal microbiome (Supplementary Figure 1A and B). Despite this, relative abundance comparisons of individual microbes at the ASV level showed increased abundance of *Bacteroides vulgatus*, *Holdemanella biformis* and *Lachnospiraceae* spp. in the fecal microbiome. The salivary microbiome showed a significant decrease (*P*<0.05) in the abundance of multiple microbes (e.g., *Prevotella histicola, Megasphaera micronuciformis, Prevotella pallens, Campylobacter concisus, Dialister invisus etc*.) in the AxSpA patients as compared with HCs (Supplementary Figure 1F).

**Figure 1.**
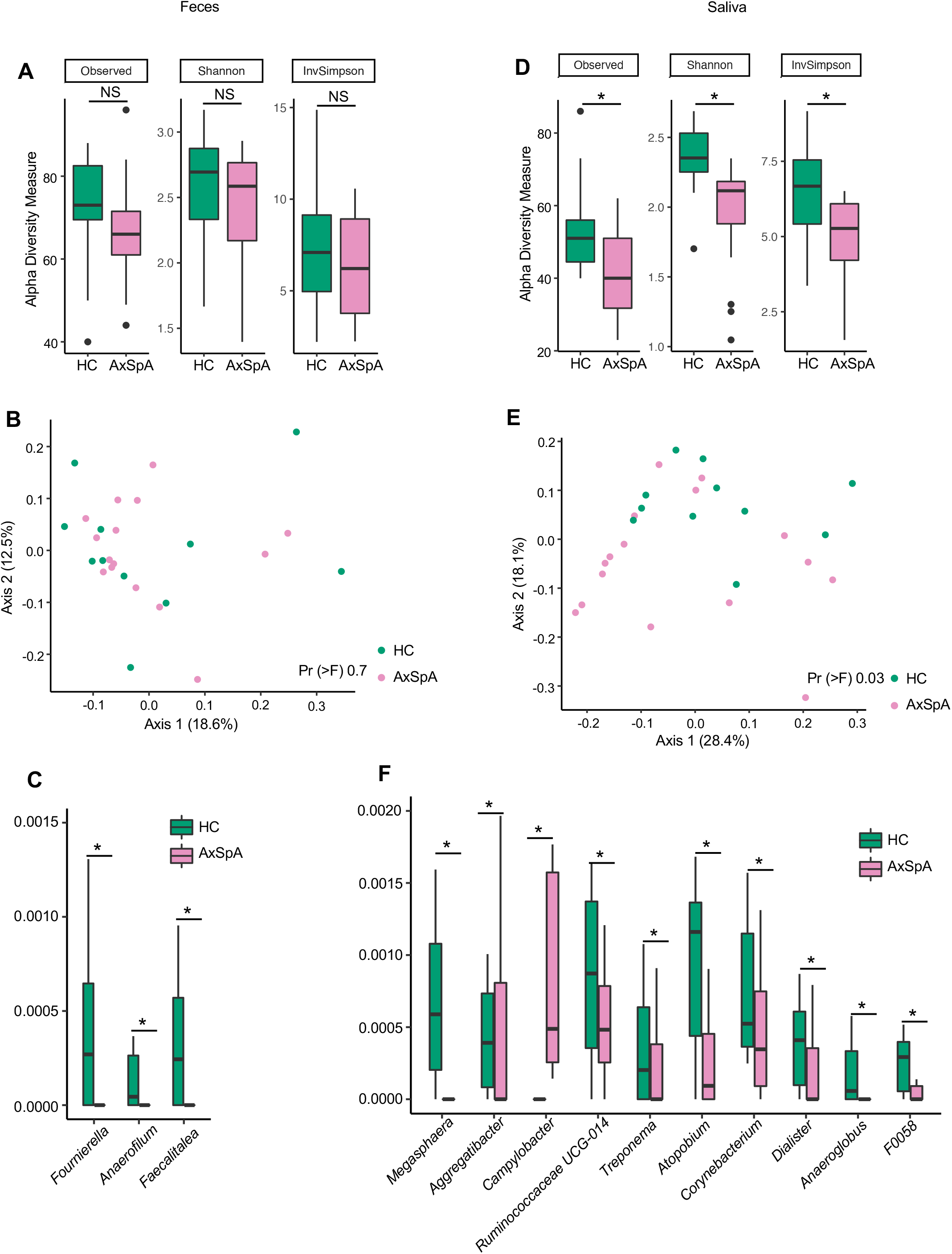
16S sequencing of fecal and salivary samples in AxSpA patients and HCs at the genus level. A-C represent fecal samples and D-F are salivary samples from AxSpA patients (pink) and HC (green). Alpha diversity plots with Observed, Shannon and Inverse Simpson (InvSimpson) indices for (A) fecal and (D) salivary samples (\**P*<0.05). Microbial composition of (B) fecal and (E) salivary samples analyzed using the unweighted Unifrac distance represented as a Principal coordinate analysis (PCoA) plot. The salivary microbial community in AxSpA patients is different from the HC (Pr (>F) 0.03). Relative abundance of genus level microbes in (C) fecal and (F) salivary samples from AxSpA patients and HC (\**P*<0.05; NS not significant).

### Enrichment of IgA-coated microbes in fecal samples of AxSpA patients

Although we did not observe differences in the gross microbial composition in fecal samples as measured by 16S sequencing, we suspected that there may be differences in how the host immune system interacted with these microbes. To determine whether microbiota from the fecal and saliva samples can elicit an immune response, we performed IgA-SEQ on IgA^+^ and IgA^−^ microbes sorted from saliva and fecal microbial communities of the AxSpA patients and HCs. This analysis revealed a trend towards increased microbial diversity of IgA^+^ fraction in the fecal microbiome of AxSpA patients as compared to the HCs, whereas the IgA^−^ fraction was less diverse in AxSpA patients as compared with HCs (Figure 2A). On the other hand, salivary microbiome of AxSpA patients showed a significant decrease in both IgA^−^ as well as IgA^+^ fractions as compared to the HCs (Figure 2E). The beta diversity comparison (IgA^+^ and IgA^−^ fractions) between AxSpA and HCs did not yield any significant differences (data not shown).

**Figure 2.**
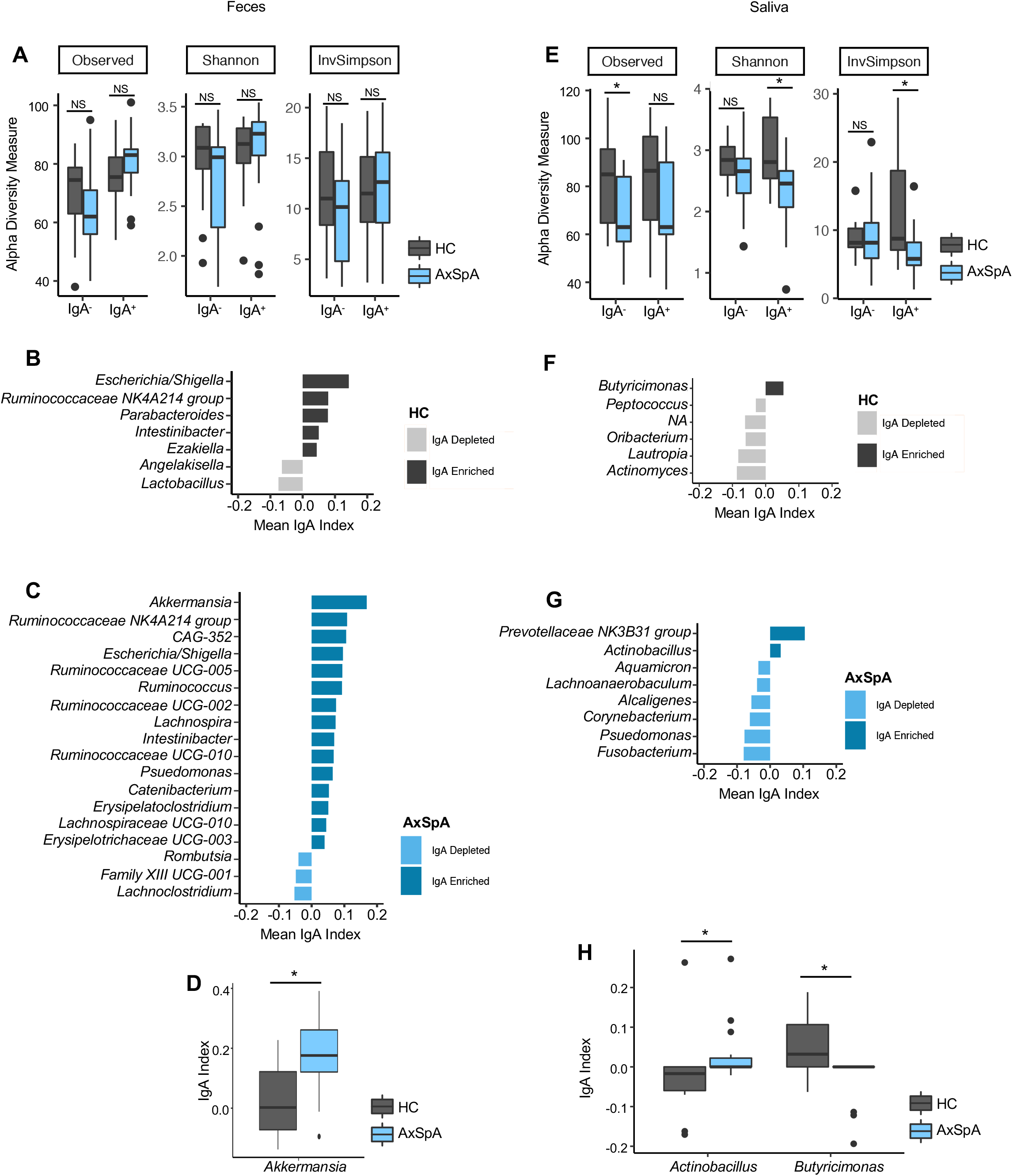
IgA-SEQ analyses of fecal and salivary samples in AxSpA patients and healthy controls (HC) at the genus level. A-D represent fecal samples and E-H are salivary samples from AxSpA patients (blue) and HC (gray). Alpha diversity plots for IgA^−^ and IgA^+^ fractions with observed, Shannon and Inverse Simpson (InvSimpson) indices for (A) fecal and (E) salivary samples (\**P*<0.05). C-D and F-G. IgA coating index was calculated for all taxa and significant taxa are shown for fecal samples for HCs (B) and patients with AxSpA (C), as well as for salivary samples from HCs (F) and AxSpA patients (G). IgA index for individual fecal (D) and salivary (H) microbes in AxSpA patients and HCs are shown (\**P*<0.05; NS not significant).

Further analyses of IgA-enriched and depleted bacteria from patients with AxSpA identified a greater number of unique IgA-coated microbes in fecal and salivary specimens at a genus (Figure 2C and 2G, respectively) and species level (Supplementary Figure 2C and 2G, respectively). *Akkermansia* has the highest IgA index score among fecal microbes in AxSpA patients followed by *Ruminococcaceae* and CAG-352. In addition, AxSpA patients had many microbes with IgA enrichment such as *Klebsiella, Ruminococcus, Lachnospira, Pseudomonas* and other members of family *Ruminococcaceae* and *Lachnospiraceae* (Figure 2C). Among fecal samples, the phylum Firmicutes represent the majority of microbes coated with IgA in AxSpA patients. The IgA^+^ fractions in HCs (Figure 2B) also showed some microbes with increased IgA coating in the fecal samples (*Escherichia/Shigella, Ruminococcaceae*-NK4A214, *Parabacteroides*). Interestingly, some of the IgA enriched microbes overlapped between AxSpA patients and HCs, such as those belonging to genus *Escherichia/Shigella, Ruminococcaceae*-NK4A214, *Intestinibacter* (Figure 2B and C). Despite significant changes in microbial diversity and community structure in the saliva of AxSpA patients in comparison to HCs (Figure 1B, D), not many microbes were altered in their IgA enrichment (Figure 2D). The majority of salivary microbes from AxSpA patients and HCs were IgA-depleted, with only the genus *Butyricimonas* being enriched in IgA coating in HCs and members of family *Prevotellaceae*, NK3831 group and *Alloprevotella* being enriched in AxSpA patients. Comparing the IgA index for individual microbes, *Akkermansia* was significantly enriched in the IgA^+^ fraction in AxSpA patients (Figure 2D). In the salivary microbiome, IgA coating of Actinobacteria was increased while that of *Butyricimonas* was decreased when compared to HCs (Figure 2H). These analyses were also performed at the ASV level and similar trends were observed (Supplementary Figure 2 A-G). Some of the IgA enriched fecal microbes such as members of *Lachnospiraceae* and *Escherichia/Shigella* overlapped between AxSpA patients and HCs (Supplementary Figures 2B and C). However, there was no overlap between the IgA enriched microbes between AxSpA patients and HCs (Supplementary Figures 2F and G).

### IgA-SEQ reveals immunoreactive microbes correlating with disease activity

To determine whether IgA enriched microbes in feces from AxSpA patients were associated with disease, we correlated genus and ASV level microbes with the disease activity score (measured by BASDAI). The analysis showed a positive correlation between IgA enrichment and disease activity only in a genus of *Clostridiales Family XIII*, and a negative correlation with the genera *Catenibacterium, Intestinibacter* and members of family *Lachnospiraceae* (UCG010) and *Ruminococcaceae* (UCG-002) (Figure 3A). At the ASV level (Figure 3B), we observed a negative correlation with (*Bacteroides* spp., *Parabacteroides johnsonii, Intestinibacter bartelettii, Firimicutes spp., Ruminococacceae spp*.). Interestingly, we did not observe disease associations with the microbes from the IgA^−^ fractions, suggesting an active role of IgA enriched microbes in disease. We did not observe significant correlations with either IgA^+^ or IgA^−^ salivary fraction with disease activity (data not shown).

**Figure 3.**
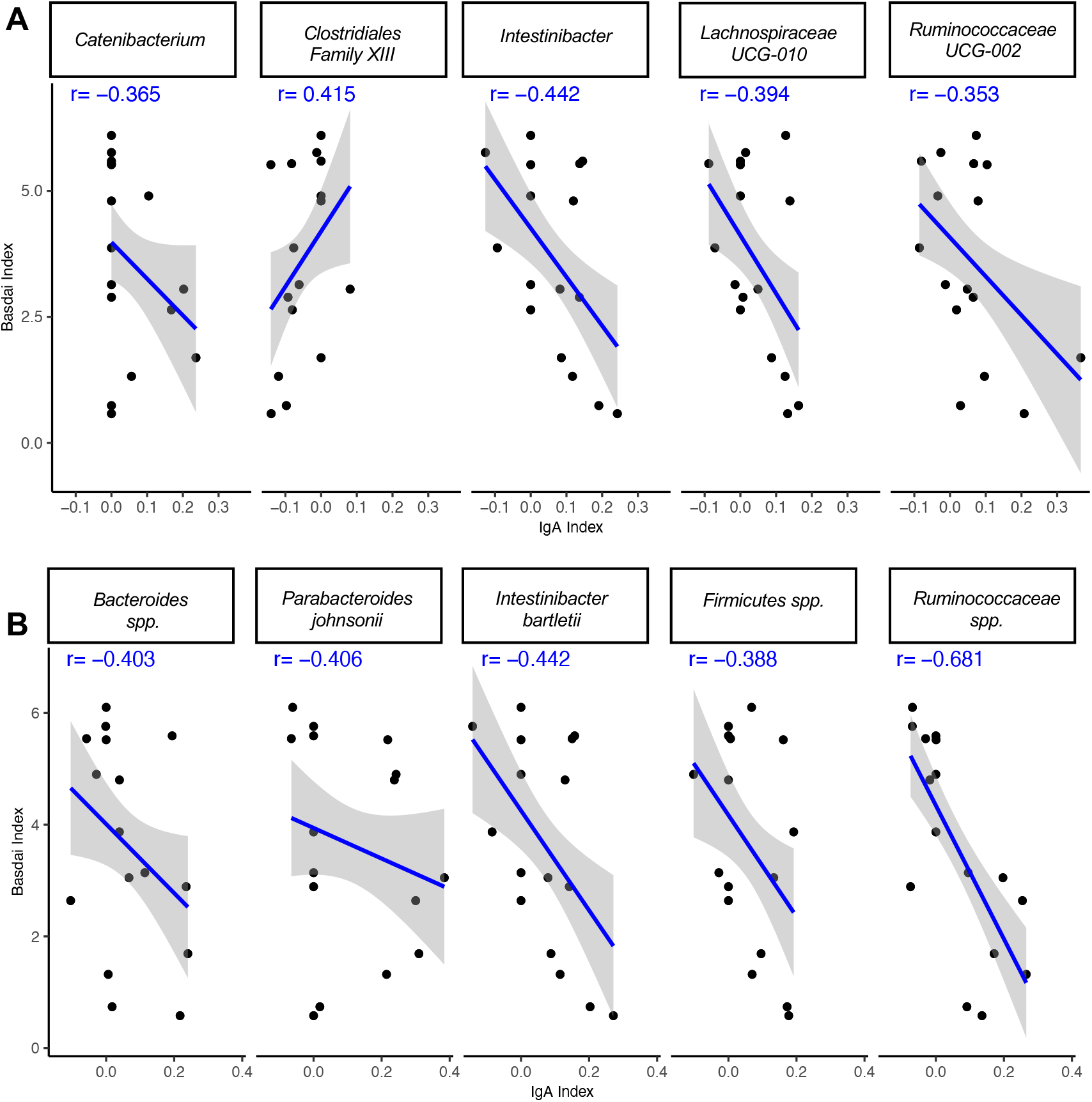
Correlation of immune reactive microbes with disease activity. The IgA index of the immune reactive microbes is correlated with the disease index (BASDAI score) at the genus (A) and ASV (B) level. The strength of each linear association is measured with a correlation coefficient r. The value of r=1 mean perfect correlation; a positive value denotes a positive correlation and a negative value signifies an inverse correlation.

### Perturbation in microbial metabolites in AxSpA patients and healthy subjects

To infer the microbial metagenome and metabolic pathways, we employed PICRUSt2, which uses marker gene data (16S gene) to predict microbial metabolic potential. We found multiple metabolic pathways (KEGG Level 3) significantly increased in the fecal IgA^+^ fraction of AxSpA patients (Figure 4A) including propanoate metabolism, bacterial secretion, N-glycan synthesis, with a decrease in metabolic pathways in the IgA^−^ fractions (Figure 4B). On the contrary, salivary IgA^+^ and IgA^−^ fractions (Figures 4C and 4D) showed increased metabolic pathways in HCs as compared with AxSpA patients. In addition, LEfSE analysis at the KEGG ortholog (KO) level for predictive genes/metabolite revealed perturbation of 112 predictive genes in the IgA^+^ fraction and 24 genes in the IgA^−^ fraction of AxSpA fecal samples (Figure 4E). Meanwhile, IgA^+^ and IgA^−^ fractions had higher numbers of predictive genes (521 and 122 respectively) perturbed in AxSpA patients, with 51 shared genes between the IgA^+^ and IgA^−^ fractions (Figure 4F). However, minimal overlap of the oral and fecal microbial genes was observed (data not shown). In the fecal IgA^+^ fraction, genes from inflammatory pathways such as lipopolysaccharide biosynthesis (K19353, K0323), glutathione metabolism (K00036, K00033, K00432), oxidative phosphorylation (K00239), biofilm formation (K01791, K00640) were perturbed. The IgA^−^ fraction showed a decreased abundance of predicted genes from metabolic pathways such as propanoate and (K01035, K01034, K00140), butanoate metabolism (K01035, K01034, K14534) in AxSpA patients. The salivary IgA^+^ fraction from AxSpA patients had decreased abundance of genes from butanoate (K01907, K17865, K03821, K00823, K01692, K01028, K07246, K01640), and propanoate (K19745, K01965, K01908, K01026, K00823, K00822, K17489) metabolism. Multiple genes for other inflammatory pathways such as tryptophan metabolism (K01692), oxidative phosphorylation (K02274, K02275, K02122, K02120, K02121, K02124, K02274, K02275, K02117, K02118), flagellar assembly (K02408, K02406, K02407), and glutathione metabolism (K07232) had altered abundances in the IgA^+^ fraction of the saliva samples. The predictive metabolites significantly altered in the IgA^+^ and IgA^−^ fractions from fecal and salivary samples are detailed in Supplementary Table S1.

**Figure 4.**
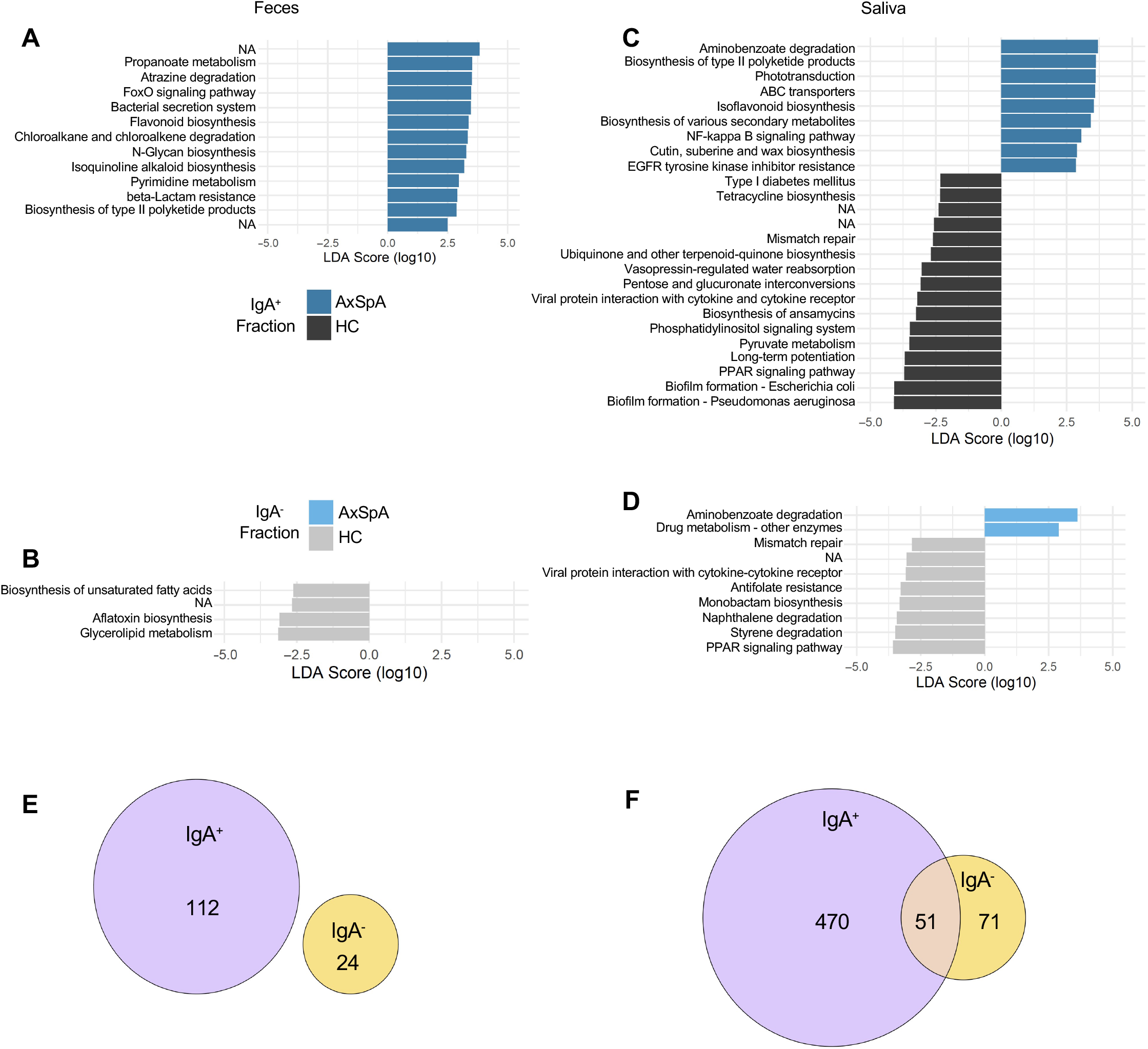
PICRUSt2 KEGG metabolites enriched in AxSpA patients and healthy individuals. Linear Discriminant analysis (LDA) effect size analysis showing differentially abundant KEGG level 3 metabolic pathways between various groups shown in a bar plot in (A) IgA enriched and (B) IgA depleted fecal fractions comparing AxSpA patients with HCs. The bar plots for salivary (C) IgA enriched and (D) IgA depleted fractions from AxSpA patients in comparison with HCs are also shown. Predicted microbial metabolites (KEGG level D) in the (E) feces and (F) saliva. The data for both IgA^+^ and IgA^−^ fractions and their overlap as shown in area proportional Euler plots. Briefly, the numbers represent KEGG genes significantly increased in their LDA score in AxSpA patients (purple) as compared with HCs (yellow). KEGG metabolites with a p<0.05 and FDR q<0.01 are considered significant.

## DISCUSSION

In this study, we examined the IgA coating of microbes in the fecal and salivary microbiome from AxSpA patients and HCs. To our knowledge, this is the first study to focus on IgA-enriched and depleted microbes and their predictive metabolic contributions in AxSpA patients. We found an enrichment of IgA coated microbes in the fecal samples from AxSpA patients. Also, the salivary microbiome from AxSpA patients had a lower number of IgA coated microbes despite showing significant microbial dysbiosis. Some of these IgA coated microbes showed an inverse correlation with the BASDAI disease activity index. In addition, predictive microbial functional revealed various inflammatory and metabolic pathways perturbed in fecal and salivary samples from AxSpA patients. Taken together, our data suggest that IgA selectively marks inflammatory or immune reactive microbes, resulting in an altered microbial community structure and function in AxSpA patients.

The present study combining 16S sequencing of the oral and fecal microbiome and the IgA coated microbes proved useful in deciphering whether dysbiotic microbes are differentially coated with IgA. Despite minor changes in the overall microbial diversity and composition in AxSpA patients and HCs, we observed a significant change in the IgA coating of oral and fecal microbes in these patients as compared to HCs. Notably, IgA-SEQ revealed many microbes that were not differentially abundant using the 16S sequencing alone, e.g., *Akkermansia*, and members of family *Ruminococcaceae* and *Lachnospiraceae*. These microbes have been shown to be associated with clinical studies in SpA and IBD (12, 31, 32) as well as in experimental models of SpA (22, 33, 34). Interestingly, *Akkermansia* has been implicated as a pathogenic microbe in SpA and IBD (12, 31, 32); and a protective microbe in metabolic diseases and obesity studies (35, 36). We also observed other IgA coated microbes in the fecal (e.g., *Lachnospiraceae, Ruminococcaceae*) and salivary (e.g., *Prevotellaceae, Actinobacillus*) samples from AxSpA patients, which have been shown to be associated with various spondyloarthropathies (12, 14). Therefore, our study highlights *Akkermansia*, and members of *Lachnospiraceae, Ruminococcaceae and Prevotellaceae* for eliciting an IgA response which suggests immunologic significance. Since these organisms are able to activate the host immune response as indicated by an IgA response, the targeted micro-organisms should be studied further for functional analysis to determine hostmicrobe interactions in disease development.

Recent studies have focused on specific microbes as a disease biomarker, patients with AS and SpA have shown disease correlation with increased relative abundance of *Ruminococcus gnavus* and *Dialister*, respectively in two different studies (12, 13). However, there are no markers for disease activity. Our study correlating various IgA coated microbes and disease activity (BASDAI) revealed many microbes with negative correlation between IgA coating and BASDAI score. However, a member of class *Clostridiales Family XIII* showed positive correlation between IgA coating and increased BASDAI score, without a significant alteration in its relative abundance in AxSpA patients (data not shown). While IgA coated microbes have shown to exacerbate gut inflammation (18, 37), IgA coating also regulates microbial composition (38). In contrast, IgA coated microbes from healthy individuals have been shown to protect from gut inflammation (39). These opposing roles for IgA may be due to the T cell dependent IgA production affected by Th-17 cells or Regulatory T cells (37, 39). These studies suggest a potential for using IgA coating as a microbial biomarker for disease activity.

Our results demonstrate alterations in the salivary microbial community associated with AxSpA patients. While the salivary microbes in AxSpA patients showed significant alterations in the microbial community and structure, we observed a smaller number of IgA coated microbes in comparison with the fecal samples. In addition, the correlation analysis between salivary microbes from the IgA^+^ and IgA^−^ fractions and the disease score (BASDAI Index) were not significant. However, there are significant changes in the predictive microbial function in both IgA^+^ and IgA^−^ fractions from saliva samples of AxSpA patients. A possible explanation is that perturbation of oral mucosa may be a result of disease instead of driving the disease in AxSpA patients.

To evaluate the functional potential of IgA enriched microbes in AxSpA patients, we employed PICRUSt2 to predict microbial metabolites altered in IgA^+^ and IgA^−^ fractions in feces and saliva. We found that most microbial metabolites perturbed in AxSpA patients belong to inflammatory pathways (e.g., biosynthesis of secondary metabolites, oxidative phosphorylation, glutathione and tryptophan metabolism, bacterial flagellar and chemotaxis genes) (40, 41)and pathways for short chain fatty acid (SCFA) metabolism (e.g., butanoate and propanoate metabolism) (34, 40, 42). Recent studies from Berlinberg and colleagues also show increased gene abundance for various metabolites in the tryptophan pathways in patients with AxSpA and AxSpA-CD (31). Alterations in tryptophan pathway are also reported by Stoll and colleagues to be associated with pediatric SpA (43). Metabolomic analysis of the plasma samples from AS patients also found a decrease in tryptophan metabolites (44). These studies indicate that decrease in tryptophan or increase in tryptophan metabolism might play an important role in AS pathogenesis. Also, multiple genes belonging to SCFA metabolism were perturbed in the saliva and fecal microbiome in AxSpA patients. Previous studies by Shao and colleagues (45) have shown that AS and RA patients have decreased production of SCFAs - butanoate and propanoate. In addition, Asquith and colleagues performed fecal metabolomic analysis in HLA-B27 transgenic rats and found many inflammatory and SCFA metabolism pathways were altered in these animals (46). Gut inflammation in these HLA-B27 transgenic rats was ameliorated by adding propanoate in their drinking water (46), further emphasizing the protective role of SCFAs. These studies suggest the need for an in-depth analysis of fecal and blood metabolites and their association with AS. Furthermore, metagenomic analysis of the IgA coated microbes will allow studying host-microbe interactions at the species or strain level.

Our study has some limitations. First, we did not observe loss of fecal microbial diversity in AxSpA patients in comparison with HCs. This may be indicative of AxSpA being a true dysbiotic model, instead of a being associated with loss in microbial diversity as reported in IBD (47, 48). Similarly, a large cohort of patients with juvenile idiopathic arthritis (JIA) (75 JIA patients and 32 controls) did not find significant differences in the alpha or beta diversity of gut microbes when compared to controls (49), despite having significant changes at the level of individual microbes. Another study performed metagenomic sequencing on fecal samples from patients with CD, AxSpA, AxSpA-CD and healthy controls did not observe any differences in the microbial alpha diversity between these groups (31). Second, our observations were limited to fecal samples, and a more severe microbial dysbiosis may be present in the intestinal mucosal microbiome as shown in our previous study in HLA-B27 TG rats (22).

The etiology of AxSpA is complex involving an interplay of genetic, environmental and microbial factors in disease development and severity (3, 50). Our results highlight the value of identifying the IgA coated microbes, which are immune-targeted microbes, based on 16S sequencing of the IgA^+^ and IgA^−^ fractions. Further in-depth examination of IgA coated microbes using shotgun metagenomic sequencing of IgA^+^ and IgA^−^ fractions to determine potentially inflammatory microbes at a strain specific level is warranted. Our results suggest that IgA coated microbes in AxSpA patients indicate a stimulated immune response to gut commensal microbes, which might contribute to the development of AxSpA. Future studies to determine the mechanistic links between IgA coated microbiota and disease will further highlight the functional implications of IgA coated bacteria in AxSpA.

## Acknowledgements

The authors wish to thank Lindsey Watson and Puthyda Keath for human subjects’ recruitment and sample processing, Manny Rodriguez for laboratory assistance, and the Spondylitis Association of America for allowing us to advertise the study to its members.

## Author Contributions

All the authors were involved in drafting or critical revisions of the manuscript and approve of the final version. Drs. Gill, Martin, Rosenbaum and Karstens had access to all the data and patient records and share the responsibly for data integrity and its accurate analysis.

---

**Supplementary Figure 1.**
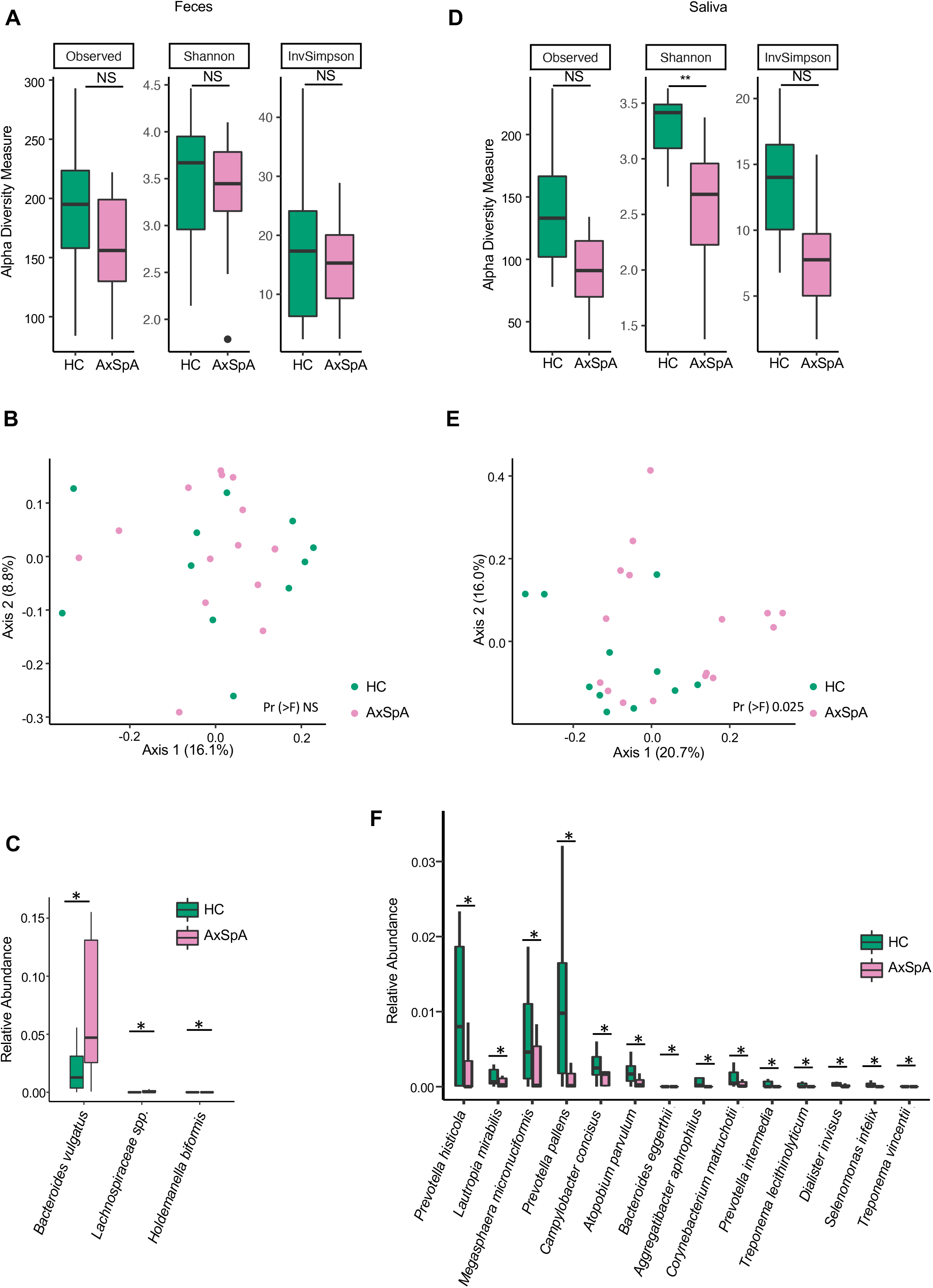
16S-sequencing of fecal and salivary samples in AxSpA patients and healthy controls (HC) at the ASV level. A-C represent fecal samples and D-F are salivary samples from AxSpA patients (pink) and HC (green). Alpha diversity plots show differences in microbial diversity at the ASV level using Observed, Shannon and Inverse Simpson (InvSimpson) indices for (A) fecal and (D) salivary samples (\**P*<0.05). Microbial composition of (B) fecal and (E) salivary samples analyzed at the ASV level using unweighted Unifrac distance. The salivary microbial community in AxSpA patients is different from the HC (Pr (>F) 0.03). Relative abundance of genus level microbes in (C) fecal and (F) salivary samples from AxSpA patients and HC (\**P*<0.05).

**Supplementary Figure 2.**
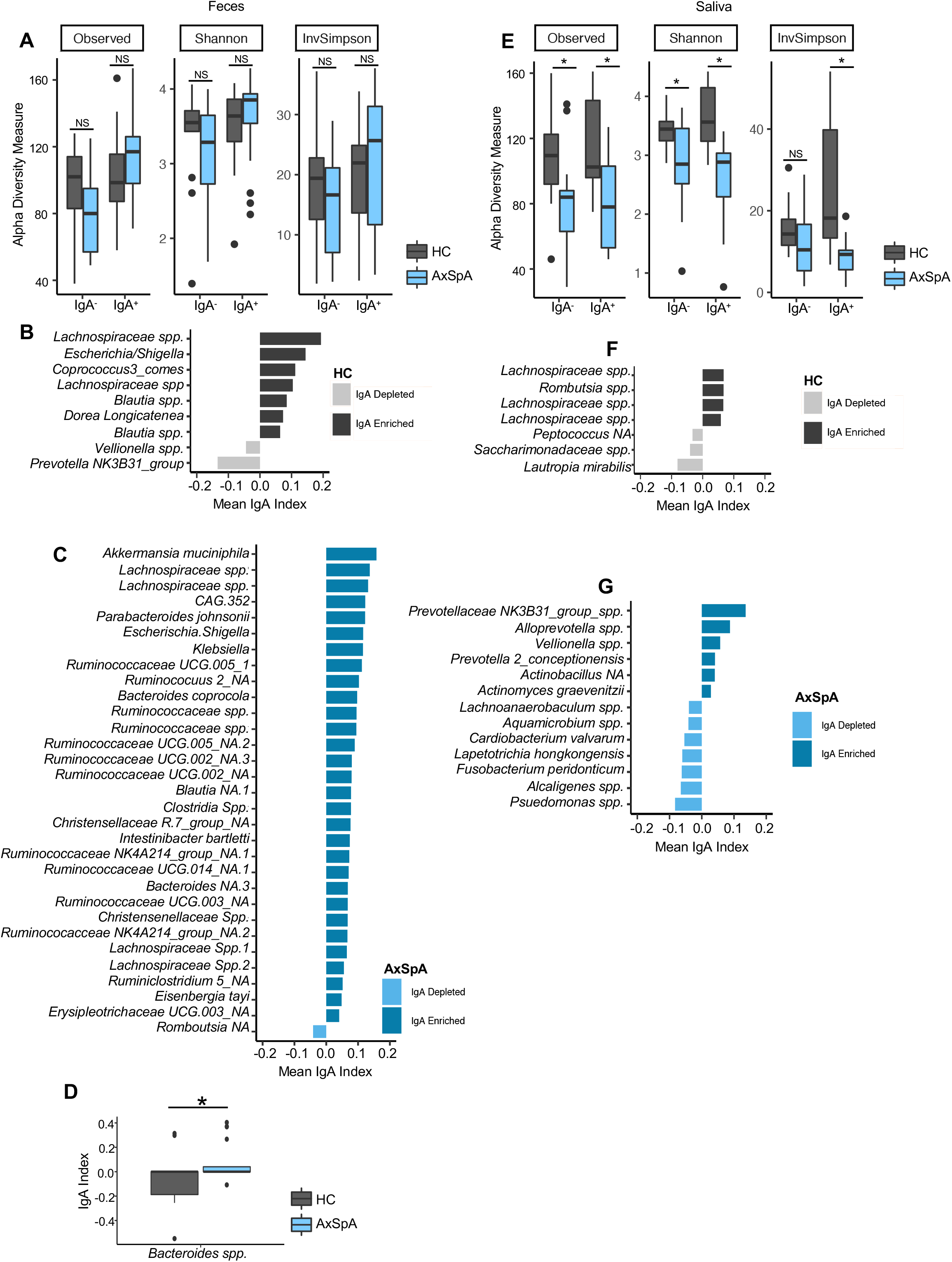
IgA-SEQ analyses of fecal and salivary samples in AxSpA patients and healthy controls (HC) at ASV level. A-D represent fecal samples and E-G are salivary samples from AxSpA patients (blue) and HC (gray). Alpha diversity plots for IgA^−^ and IgA^+^ fractions with Observed, Shannon and Inverse Simpson (InvSimpson) indices for (A) fecal and (E) salivary samples (\**P*<0.05). C-D and F-G. IgA coating index was calculated for all taxa and significant taxa () are shown for fecal samples for HCs (B) and patients with AxSpA (C), as well as for salivary samples from HCs (F) and AxSpA patients (G). IgA index for individual fecal (D) microbes in AxSpA patients and HCs are shown (\**P*<0.05).

**Supplementary Table S1**. Predictive microbial metabolites (KEGG level 4) significantly altered in AxSpA patients in comparison with HCs from IgA^+^ and IgA^−^ fractions of fecal and salivary samples.

