## Supplemental Table 1 for "Axial spondyloarthritis patients have altered mucosal IgA response to oral and fecal microbiota"

**Supplementary Table S1**. Predictive microbial metabolites (KEGG level 4) significantly altered in AxSpA patients in comparison with HCs from IgA+ and IgA- fractions of feces and saliva samples.

| S_ASvHC_IgA-_a0.05_w0.05_l2_minc.default.res.sig | | | | |
| --- | --- | --- | --- | --- |
| LevelD_(KO) Description | KO | Class with highest mean | Log LDA score | p-value (KW for class) |
| SCD, desC; stearoyl-CoA desaturase (Delta-9 desaturase) [EC:1.14.19.1] | K00507 | HC | 2.66 | 0.01 |
| phoH2; PhoH-like ATPase | K07175 | HC | 2.67 | 0.01 |
| ssuB; sulfonate transport system ATP-binding protein [EC:7.6.2.14] | K15555 | AxSpA | 2.85 | 0.04 |
| LYSN; 2-aminoadipate transaminase [EC:2.6.1.-] | K05825 | HC | 3.47 | 0.01 |
| xdhC; xanthine dehydrogenase accessory factor | K07402 | HC | 3.81 | 0.02 |
| mcp; methyl-accepting chemotaxis protein | K03406 | HC | 3.85 | 0.04 |
| cheA; two-component system, chemotaxis family, sensor kinase CheA [EC:2.7.13.3] | K03407 | HC | 3.02 | 0.04 |
| cheW; purine-binding chemotaxis protein CheW | K03408 | HC | 3.10 | 0.03 |
| K07099; uncharacterized protein | K07099 | HC | 3.42 | 0.03 |
| tagA, tarA; N-acetylglucosaminyldiphosphoundecaprenol N-acetyl-beta-D-mannosaminyltransferase [EC:2.4.1.187] | K05946 | HC | 3.34 | 0.04 |
| hisI; phosphoribosyl-AMP cyclohydrolase [EC:3.5.4.19] | K01496 | HC | 3.28 | 0.03 |
| fliO, fliZ; flagellar protein FliO/FliZ | K02418 | HC | 2.92 | 0.03 |
| fliJ; flagellar protein FliJ | K02413 | HC | 2.93 | 0.02 |
| fliN; flagellar motor switch protein FliN | K02417 | HC | 3.05 | 0.03 |
| fliK; flagellar hook-length control protein FliK | K02414 | HC | 2.80 | 0.01 |
| agaR; DeoR family transcriptional regulator, aga operon transcriptional repressor | K02081 | HC | 2.59 | 0.01 |
| addA; ATP-dependent helicase/nuclease subunit A [EC:3.1.-.- 3.6.4.12] | K16898 | HC | 3.47 | 0.05 |
| MUT; methylmalonyl-CoA mutase [EC:5.4.99.2] | K01847 | HC | 3.58 | 0.04 |
| dinD; DNA-damage-inducible protein D | K14623 | HC | 3.23 | 0.04 |
| PCBD, phhB; 4a-hydroxytetrahydrobiopterin dehydratase [EC:4.2.1.96] | K01724 | HC | 2.63 | 0.02 |
| fliB; lysine-N-methylase [EC:2.1.1.-] | K18475 | HC | 3.44 | 0.03 |
| ELP3, KAT9; elongator complex protein 3 [EC:2.3.1.48] | K07739 | HC | 2.44 | 0.00 |
| paaI; acyl-CoA thioesterase [EC:3.1.2.-] | K02614 | HC | 3.33 | 0.04 |
| imuA; protein ImuA | K14160 | HC | 2.67 | 0.02 |
| larE; pyridinium-3,5-biscarboxylic acid mononucleotide sulfurtransferase [EC:4.4.1.37] | K06864 | HC | 3.30 | 0.03 |
| flgM; negative regulator of flagellin synthesis FlgM | K02398 | HC | 2.94 | 0.03 |
| flgJ; peptidoglycan hydrolase FlgJ | K02395 | HC | 2.73 | 0.00 |
| flgL; flagellar hook-associated protein 3 FlgL | K02397 | HC | 2.93 | 0.02 |
| flgE; flagellar hook protein FlgE | K02390 | HC | 3.04 | 0.03 |
| yabN; tetrapyrrole methylase family protein / MazG family protein | K02499 | HC | 3.42 | 0.03 |
| hoxH; NAD-reducing hydrogenase large subunit [EC:1.12.1.2] | K00436 | HC | 2.81 | 0.02 |
| E2.1.3.1-1.3S; methylmalonyl-CoA carboxyltransferase 1.3S subunit [EC:2.1.3.1] | K17490 | HC | 2.48 | 0.04 |
| E2.6.1.83; LL-diaminopimelate aminotransferase [EC:2.6.1.83] | K10206 | HC | 3.58 | 0.05 |
| ABC.MR; putative ABC transport system ATP-binding protein | K02021 | HC | 2.36 | 0.05 |
| rfbF, rhlC; rhamnosyltransferase [EC:2.4.1.-] | K12990 | HC | 2.46 | 0.01 |
| mcsA; protein arginine kinase activator | K19411 | HC | 3.34 | 0.04 |
| fbaB; fructose-bisphosphate aldolase, class I [EC:4.1.2.13] | K11645 | HC | 2.75 | 0.01 |
| cpaB, rcpC; pilus assembly protein CpaB | K02279 | HC | 2.68 | 0.00 |
| E3.3.1.1, ahcY; adenosylhomocysteinase [EC:3.3.1.1] | K01251 | HC | 3.42 | 0.01 |
| hisZ; ATP phosphoribosyltransferase regulatory subunit | K02502 | HC | 3.51 | 0.04 |
| K09859; uncharacterized protein | K09859 | HC | 2.38 | 0.02 |
| E3.2.1.58; glucan 1,3-beta-glucosidase [EC:3.2.1.58] | K01210 | HC | 2.34 | 0.00 |
| croR; 3-hydroxybutyryl-CoA dehydratase [EC:4.2.1.55] | K17865 | HC | 2.77 | 0.01 |
| nox2; NADH oxidase (H2O-forming) [EC:1.6.3.4] | K17869 | HC | 2.41 | 0.00 |
| pct; propionate CoA-transferase [EC:2.8.3.1] | K01026 | HC | 2.75 | 0.04 |
| fliH; flagellar assembly protein FliH | K02411 | HC | 2.89 | 0.02 |
| dfx; superoxide reductase [EC:1.15.1.2] | K05919 | HC | 3.52 | 0.03 |
| cfa; cyclopropane-fatty-acyl-phospholipid synthase [EC:2.1.1.79] | K00574 | HC | 3.34 | 0.03 |
| nifH; nitrogenase iron protein NifH | K02588 | HC | 2.81 | 0.05 |
| nifE; nitrogenase molybdenum-cofactor synthesis protein NifE | K02587 | HC | 2.54 | 0.04 |
| PITRM1, PreP, CYM1; presequence protease [EC:3.4.24.-] | K06972 | HC | 3.40 | 0.04 |
| K07089; uncharacterized protein | K07089 | HC | 3.43 | 0.05 |
| dat; D-alanine transaminase [EC:2.6.1.21] | K00824 | HC | 3.31 | 0.02 |
| fliD; flagellar hook-associated protein 2 | K02407 | HC | 3.05 | 0.02 |
| fliC, hag; flagellin | K02406 | HC | 3.30 | 0.04 |
| mocA; molybdenum cofactor cytidylyltransferase [EC:2.7.7.76] | K07141 | HC | 3.40 | 0.01 |
| E4.2.1.2AA, fumA; fumarate hydratase subunit alpha [EC:4.2.1.2] | K01677 | HC | 3.51 | 0.02 |
| E4.2.1.2AB, fumB; fumarate hydratase subunit beta [EC:4.2.1.2] | K01678 | HC | 3.51 | 0.02 |
| cheB; two-component system, chemotaxis family, protein-glutamate methylesterase/glutaminase [EC:3.1.1.61 3.5.1.44] | K03412 | HC | 2.98 | 0.03 |
| E2.1.3.1-5S; methylmalonyl-CoA carboxyltransferase 5S subunit [EC:2.1.3.1] | K03416 | HC | 2.48 | 0.04 |
| hemDX; uroporphyrinogen III methyltransferase / synthase [EC:2.1.1.107 4.2.1.75] | K13543 | HC | 2.59 | 0.04 |
| K07317; adenine-specific DNA-methyltransferase [EC:2.1.1.72] | K07317 | HC | 2.13 | 0.03 |
| spoVB; stage V sporulation protein B | K06409 | HC | 3.72 | 0.04 |
| larC; pyridinium-3,5-bisthiocarboxylic acid mononucleotide nickel chelatase [EC:4.99.1.12] | K09121 | HC | 3.35 | 0.02 |
| fliF; flagellar M-ring protein FliF | K02409 | HC | 2.95 | 0.03 |
| hisB; imidazoleglycerol-phosphate dehydratase [EC:4.2.1.19] | K01693 | HC | 3.50 | 0.02 |
| livM; branched-chain amino acid transport system permease protein | K01998 | HC | 3.67 | 0.01 |
| livK; branched-chain amino acid transport system substrate-binding protein | K01999 | HC | 3.74 | 0.04 |
| livF; branched-chain amino acid transport system ATP-binding protein | K01996 | HC | 3.66 | 0.01 |
| livH; branched-chain amino acid transport system permease protein | K01997 | HC | 3.56 | 0.01 |
| livG; branched-chain amino acid transport system ATP-binding protein | K01995 | HC | 3.62 | 0.01 |
| treS; maltose alpha-D-glucosyltransferase / alpha-amylase [EC:5.4.99.16 3.2.1.1] | K05343 | HC | 2.71 | 0.02 |
| K09726; uncharacterized protein | K09726 | HC | 2.63 | 0.01 |
| glcD; glycolate oxidase [EC:1.1.3.15] | K00104 | HC | 3.18 | 0.01 |
| cobL-cbiET; precorrin-6B C5,15-methyltransferase / cobalt-precorrin-6B C5,C15-methyltransferase [EC:2.1.1.132 2.1.1.289 2.1.1.196] | K00595 | HC | 3.18 | 0.02 |
| HK; hexokinase [EC:2.7.1.1] | K00844 | HC | 2.66 | 0.05 |
| K06876; deoxyribodipyrimidine photolyase-related protein | K06876 | HC | 2.91 | 0.03 |
| hisE; phosphoribosyl-ATP pyrophosphohydrolase [EC:3.6.1.31] | K01523 | HC | 3.27 | 0.03 |
| flgD; flagellar basal-body rod modification protein FlgD | K02389 | HC | 2.94 | 0.03 |
| K09807; uncharacterized protein | K09807 | HC | 3.33 | 0.04 |
| pfkB; 6-phosphofructokinase 2 [EC:2.7.1.11] | K16370 | HC | 2.15 | 0.01 |
| hpdB; 4-hydroxyphenylacetate decarboxylase large subunit [EC:4.1.1.83] | K18427 | HC | 2.30 | 0.00 |
| ybiV; sugar-phosphatase [EC:3.1.3.23] | K07757 | HC | 2.24 | 0.01 |
| cobC1, cobC; cobalamin biosynthesis protein CobC | K02225 | HC | 2.59 | 0.03 |
| cobF; precorrin-6A synthase [EC:2.1.1.152] | K02228 | HC | 2.44 | 0.04 |
| mtnA; methylthioribose-1-phosphate isomerase [EC:5.3.1.23] | K08963 | HC | 3.33 | 0.04 |
| torR; two-component system, OmpR family, torCAD operon response regulator TorR | K07772 | HC | 2.82 | 0.03 |
| ccr; crotonyl-CoA carboxylase/reductase [EC:1.3.1.85] | K14446 | HC | 2.78 | 0.04 |
| dgoT; MFS transporter, ACS family, D-galactonate transporter | K08194 | HC | 2.37 | 0.03 |
| cdgJ; c-di-GMP phosphodiesterase [EC:3.1.4.52] | K07181 | HC | 2.67 | 0.02 |
| citR; LysR family transcriptional regulator, repressor for citA | K19242 | HC | 2.39 | 0.04 |
| flhB2; flagellar biosynthesis protein | K04061 | HC | 2.94 | 0.03 |
| mcsB; protein arginine kinase [EC:2.7.14.1] | K19405 | HC | 3.34 | 0.04 |
| aroG, aroA; 3-deoxy-7-phosphoheptulonate synthase / chorismate mutase [EC:2.5.1.54 5.4.99.5] | K13853 | HC | 2.33 | 0.04 |
| K07571; S1 RNA binding domain protein | K07571 | HC | 3.36 | 0.05 |
| ABC.MN.S; manganese/iron transport system substrate-binding protein | K09818 | HC | 2.24 | 0.02 |
| queE; 7-carboxy-7-deazaguanine synthase [EC:4.3.99.3] | K10026 | HC | 3.29 | 0.05 |
| pksJ; polyketide synthase PksJ | K13611 | HC | 2.33 | 0.01 |
| hydN; electron transport protein HydN | K05796 | HC | 2.15 | 0.05 |
| rihC; non-specific riboncleoside hydrolase [EC:3.2.-.-] | K12700 | HC | 2.12 | 0.02 |
| ACADM, acd; acyl-CoA dehydrogenase [EC:1.3.8.7] | K00249 | HC | 2.84 | 0.03 |
| larB; pyridinium-3,5-biscarboxylic acid mononucleotide synthase [EC:2.5.1.143] | K06898 | HC | 3.30 | 0.03 |
| ACR3, arsB; arsenite transporter | K03325 | HC | 2.72 | 0.04 |
| amt, AMT, MEP; ammonium transporter, Amt family | K03320 | HC | 3.54 | 0.02 |
| AKR1A1, adh; alcohol dehydrogenase (NADP+) [EC:1.1.1.2] | K00002 | HC | 2.58 | 0.01 |
| GLUD1_2, gdhA; glutamate dehydrogenase (NAD(P)+) [EC:1.4.1.3] | K00261 | HC | 3.29 | 0.04 |
| mtaD; 5-methylthioadenosine/S-adenosylhomocysteine deaminase [EC:3.5.4.31 3.5.4.28] | K12960 | HC | 3.38 | 0.04 |
| fosB; metallothiol transferase [EC:2.5.1.-] | K11210 | HC | 2.63 | 0.04 |
| tdh; threonine 3-dehydrogenase [EC:1.1.1.103] | K00060 | HC | 2.21 | 0.02 |
| GLU, gltS; glutamate synthase (ferredoxin) [EC:1.4.7.1] | K00284 | HC | 3.40 | 0.04 |
| cpaE, tadZ; pilus assembly protein CpaE | K02282 | HC | 2.26 | 0.00 |
| ispDF; 2-C-methyl-D-erythritol 4-phosphate cytidylyltransferase / 2-C-methyl-D-erythritol 2,4-cyclodiphosphate synthase [EC:2.7.7.60 4.6.1.12] | K12506 | HC | 3.39 | 0.02 |
| ttuC, dmlA; tartrate dehydrogenase/decarboxylase / D-malate dehydrogenase [EC:1.1.1.93 4.1.1.73 1.1.1.83] | K07246 | HC | 2.59 | 0.03 |
| TC.SMR3; small multidrug resistance family-3 protein | K09771 | HC | 3.33 | 0.04 |
| pydC; beta-ureidopropionase / N-carbamoyl-L-amino-acid hydrolase [EC:3.5.1.6 3.5.1.87] | K06016 | HC | 3.32 | 0.02 |
| pcaC; 4-carboxymuconolactone decarboxylase [EC:4.1.1.44] | K01607 | HC | 3.54 | 0.02 |
| rbcL, cbbL; ribulose-bisphosphate carboxylase large chain [EC:4.1.1.39] | K01601 | HC | 3.29 | 0.03 |
| wcaJ; putative colanic acid biosysnthesis UDP-glucose lipid carrier transferase | K03606 | HC | 2.81 | 0.02 |
| kdgT; 2-keto-3-deoxygluconate permease | K02526 | HC | 2.24 | 0.04 |
| esxA, esat6; 6 kDa early secretory antigenic target | K14956 | HC | 3.12 | 0.04 |
| remA; extracellular matrix regulatory protein A | K09777 | HC | 3.43 | 0.02 |
| thiT; thiamine transporter | K16789 | HC | 3.39 | 0.04 |

| S_ASvHC_IgA+_a0.05_w0.05_l2_minc.default.res.sig.txt | | | | |
| --- | --- | --- | --- | --- |
| LevelD (KO) Description | KO | Class with highest mean | Log LDA score | p-value (KW for class) |
| urtE; urea transport system ATP-binding protein | K11963 | HC | 2.41 | 0.01 |
| arnD; undecaprenyl phosphate-alpha-L-ara4FN deformylase [EC:3.5.1.-] | K13014 | AxSpA | 2.31 | 0.02 |
| phoH2; PhoH-like ATPase | K07175 | HC | 2.74 | 0.03 |
| bisC; biotin/methionine sulfoxide reductase [EC:1.-.-.-] | K08351 | AxSpA | 2.60 | 0.02 |
| fdnI; formate dehydrogenase-N, gamma subunit | K08350 | AxSpA | 2.28 | 0.02 |
| aaeB; p-hydroxybenzoic acid efflux pump subunit AaeB | K03468 | AxSpA | 2.28 | 0.02 |
| thyX, thy1; thymidylate synthase (FAD) [EC:2.1.1.148] | K03465 | HC | 3.19 | 0.03 |
| E4.1.3.4, HMGCL, hmgL; hydroxymethylglutaryl-CoA lyase [EC:4.1.3.4] | K01640 | HC | 2.62 | 0.00 |
| pinR; putative DNA-invertase from lambdoid prophage Rac | K14060 | HC | 2.74 | 0.01 |
| ompN; outer membrane protein N | K14062 | AxSpA | 2.35 | 0.02 |
| uspC; universal stress protein C | K14064 | AxSpA | 2.28 | 0.02 |
| solA; N-methyl-L-tryptophan oxidase [EC:1.5.3.-] | K02846 | AxSpA | 2.53 | 0.02 |
| mug; double-stranded uracil-DNA glycosylase [EC:3.2.2.28] | K03649 | AxSpA | 2.55 | 0.01 |
| trbL; type IV secretion system protein TrbL | K07344 | HC | 2.41 | 0.00 |
| K06960; uncharacterized protein | K06960 | HC | 3.46 | 0.04 |
| dauA; D-arginine dehydrogenase [EC:1.4.99.6] | K19746 | HC | 2.11 | 0.03 |
| acuI; acrylyl-CoA reductase (NADPH) [EC:1.3.1.-] | K19745 | HC | 2.13 | 0.02 |
| otsA; trehalose 6-phosphate synthase [EC:2.4.1.15 2.4.1.347] | K00697 | HC | 2.08 | 0.04 |
| lrgA; holin-like protein | K05338 | AxSpA | 2.25 | 0.02 |
| lrgB; holin-like protein LrgB | K05339 | AxSpA | 2.26 | 0.03 |
| csgD; LuxR family transcriptional regulator, csgAB operon transcriptional regulatory protein | K04333 | AxSpA | 2.28 | 0.02 |
| K02477; two-component system, LytTR family, response regulator | K02477 | HC | 3.32 | 0.01 |
| mpaA; murein peptide amidase A | K14054 | AxSpA | 2.66 | 0.02 |
| rhaS; AraC family transcriptional regulator, L-rhamnose operon regulatory protein RhaS | K02855 | AxSpA | 2.27 | 0.04 |
| sugE; quaternary ammonium compound-resistance protein SugE | K11741 | HC | 2.60 | 0.03 |
| K09928; uncharacterized protein | K09928 | HC | 2.21 | 0.01 |
| K09766; uncharacterized protein | K09766 | HC | 2.97 | 0.05 |
| K07157; uncharacterized protein | K07157 | HC | 2.30 | 0.01 |
| xylA; xylose isomerase [EC:5.3.1.5] | K01805 | HC | 2.96 | 0.02 |
| rpiA; ribose 5-phosphate isomerase A [EC:5.3.1.6] | K01807 | AxSpA | 3.77 | 0.02 |
| mcp; methyl-accepting chemotaxis protein | K03406 | HC | 4.19 | 0.03 |
| cheA; two-component system, chemotaxis family, sensor kinase CheA [EC:2.7.13.3] | K03407 | HC | 3.32 | 0.03 |
| cheW; purine-binding chemotaxis protein CheW | K03408 | HC | 3.34 | 0.04 |
| cheX; chemotaxis protein CheX | K03409 | HC | 2.56 | 0.01 |
| dnaT; DNA replication protein DnaT | K02317 | AxSpA | 2.28 | 0.02 |
| bhsA; multiple stress resistance protein BhsA | K12151 | AxSpA | 2.28 | 0.02 |
| tesA; acyl-CoA thioesterase I [EC:3.1.2.- 3.1.2.2 3.1.1.2 3.1.1.5] | K10804 | HC | 2.44 | 0.01 |
| rob; AraC family transcriptional regulator, mar-sox-rob regulon activator | K05804 | AxSpA | 2.08 | 0.03 |
| lysK; lysyl-tRNA synthetase, class I [EC:6.1.1.6] | K04566 | HC | 2.28 | 0.03 |
| flhG, fleN; flagellar biosynthesis protein FlhG | K04562 | HC | 3.04 | 0.02 |
| E2.6.1.11, argD; acetylornithine aminotransferase [EC:2.6.1.11] | K00818 | HC | 3.12 | 0.01 |
| fliO, fliZ; flagellar protein FliO/FliZ | K02418 | HC | 3.22 | 0.02 |
| fliJ; flagellar protein FliJ | K02413 | HC | 3.24 | 0.02 |
| fliG; flagellar motor switch protein FliG | K02410 | HC | 3.36 | 0.05 |
| fliK; flagellar hook-length control protein FliK | K02414 | HC | 3.10 | 0.02 |
| fliL; flagellar protein FliL | K02415 | HC | 2.46 | 0.01 |
| pdc; phenolic acid decarboxylase [EC:4.1.1.-] | K13727 | AxSpA | 2.41 | 0.02 |
| gfrF; fructoselysine-6-phosphate deglycase | K19510 | HC | 2.34 | 0.01 |
| arsH; arsenical resistance protein ArsH | K11811 | HC | 2.07 | 0.02 |
| MuB; ATP-dependent target DNA activator | K07132 | HC | 2.75 | 0.02 |
| agaR; DeoR family transcriptional regulator, aga operon transcriptional repressor | K02081 | HC | 2.77 | 0.03 |
| tatE; sec-independent protein translocase protein TatE | K03425 | AxSpA | 2.28 | 0.02 |
| lpqC; polyhydroxybutyrate depolymerase | K03932 | HC | 2.09 | 0.04 |
| E3.1.3.15B; histidinol-phosphatase (PHP family) [EC:3.1.3.15] | K04486 | HC | 3.76 | 0.03 |
| tarJ; ribitol-5-phosphate 2-dehydrogenase (NADP+) [EC:1.1.1.405] | K05352 | AxSpA | 3.16 | 0.03 |
| arsC; arsenate reductase (thioredoxin) [EC:1.20.4.4] | K03741 | HC | 3.21 | 0.04 |
| ompC; outer membrane pore protein C | K09475 | AxSpA | 2.43 | 0.02 |
| ompF; outer membrane pore protein F | K09476 | AxSpA | 2.28 | 0.02 |
| K13652; AraC family transcriptional regulator | K13652 | HC | 2.65 | 0.03 |
| ramB; XRE family transcriptional regulator, fatty acid utilization regulator | K07110 | HC | 2.16 | 0.00 |
| PTGR3, ZADH2; prostaglandin reductase 3 [EC:1.3.1.48] | K07119 | AxSpA | 2.17 | 0.02 |
| pepM; phosphoenolpyruvate phosphomutase [EC:5.4.2.9] | K01841 | HC | 2.26 | 0.01 |
| ptsP; phosphotransferase system, enzyme I, PtsP [EC:2.7.3.9] | K08484 | HC | 2.11 | 0.03 |
| mdtL; MFS transporter, DHA1 family, multidrug resistance protein | K08163 | AxSpA | 2.24 | 0.02 |
| aqpZ; aquaporin Z | K06188 | AxSpA | 3.41 | 0.01 |
| folX; D-erythro-7,8-dihydroneopterin triphosphate epimerase [EC:5.1.99.7] | K07589 | AxSpA | 2.16 | 0.04 |
| ERCC3, XPB; DNA excision repair protein ERCC-3 [EC:3.6.4.12] | K10843 | HC | 2.34 | 0.04 |
| GLPF; glycerol uptake facilitator protein | K02440 | AxSpA | 3.63 | 0.04 |
| NEU1; sialidase-1 [EC:3.2.1.18] | K01186 | AxSpA | 3.56 | 0.01 |
| cedA; cell division activator | K15722 | AxSpA | 2.28 | 0.02 |
| syd; SecY interacting protein Syd | K15723 | AxSpA | 2.29 | 0.02 |
| czcB, cusB, cnrB; membrane fusion protein, heavy metal efflux system | K15727 | HC | 2.67 | 0.01 |
| abmG; 2-aminobenzoate-CoA ligase [EC:6.2.1.32] | K08295 | AxSpA | 2.90 | 0.03 |
| PCBD, phhB; 4a-hydroxytetrahydrobiopterin dehydratase [EC:4.2.1.96] | K01724 | HC | 2.36 | 0.03 |
| pel; pectate lyase [EC:4.2.2.2] | K01728 | HC | 2.63 | 0.00 |
| uspF; universal stress protein F | K14061 | AxSpA | 2.28 | 0.02 |
| E4.1.1.82; phosphonopyruvate decarboxylase [EC:4.1.1.82] | K09459 | HC | 2.26 | 0.01 |
| K09986; uncharacterized protein | K09986 | HC | 2.15 | 0.01 |
| nfsA; nitroreductase [EC:1.-.-.-] | K10678 | HC | 2.94 | 0.02 |
| grdB; glycine reductase complex component B subunit gamma [EC:1.21.4.2] | K10672 | HC | 3.03 | 0.01 |
| yjjG; 5'-nucleotidase [EC:3.1.3.5] | K08723 | AxSpA | 2.34 | 0.02 |
| pcaH; protocatechuate 3,4-dioxygenase, beta subunit [EC:1.13.11.3] | K00449 | HC | 2.14 | 0.01 |
| cobD; threonine-phosphate decarboxylase [EC:4.1.1.81] | K04720 | HC | 3.75 | 0.02 |
| ycfS; L,D-transpeptidase YcfS | K19236 | AxSpA | 2.24 | 0.02 |
| ybiS; L,D-transpeptidase YbiS | K19235 | AxSpA | 2.22 | 0.02 |
| ABC.VB1X.S; putative thiamine transport system substrate-binding protein | K05777 | AxSpA | 2.47 | 0.05 |
| pspC; phage shock protein C | K03973 | HC | 2.67 | 0.03 |
| pspD; phage shock protein D | K03971 | AxSpA | 2.28 | 0.02 |
| ampH; serine-type D-Ala-D-Ala carboxypeptidase/endopeptidase [EC:3.4.16.4 3.4.21.-] | K18988 | AxSpA | 2.27 | 0.02 |
| psuG; pseudouridylate synthase [EC:4.2.1.70] | K16329 | HC | 3.06 | 0.01 |
| psuK; pseudouridine kinase [EC:2.7.1.83] | K16328 | HC | 3.06 | 0.00 |
| dcuS; two-component system, CitB family, sensor histidine kinase DcuS [EC:2.7.13.3] | K07701 | AxSpA | 2.43 | 0.02 |
| dpiB, citA; two-component system, CitB family, cit operon sensor histidine kinase CitA [EC:2.7.13.3] | K07700 | AxSpA | 2.01 | 0.04 |
| dcuR; two-component system, CitB family, response regulator DcuR | K07703 | AxSpA | 2.31 | 0.04 |
| dpiA, citB; two-component system, CitB family, response regulator CitB | K07702 | AxSpA | 2.05 | 0.03 |
| glnL, ntrB; two-component system, NtrC family, nitrogen regulation sensor histidine kinase GlnL [EC:2.7.13.3] | K07708 | HC | 2.31 | 0.01 |
| K09937; uncharacterized protein | K09937 | HC | 2.25 | 0.04 |
| ELP3, KAT9; elongator complex protein 3 [EC:2.3.1.48] | K07739 | HC | 2.25 | 0.01 |
| imuA; protein ImuA | K14160 | HC | 2.28 | 0.02 |
| dnaE2; error-prone DNA polymerase [EC:2.7.7.7] | K14162 | HC | 2.37 | 0.03 |
| hrcA; heat-inducible transcriptional repressor | K03705 | HC | 3.50 | 0.03 |
| matB; malonyl-CoA/methylmalonyl-CoA synthetase [EC:6.2.1.-] | K18661 | HC | 2.09 | 0.05 |
| betB, gbsA; betaine-aldehyde dehydrogenase [EC:1.2.1.8] | K00130 | HC | 2.61 | 0.03 |
| arsA, ASNA1, GET3; arsenite/tail-anchored protein-transporting ATPase [EC:7.3.2.7 7.3.-.-] | K01551 | HC | 2.40 | 0.03 |
| flgM; negative regulator of flagellin synthesis FlgM | K02398 | HC | 3.26 | 0.02 |
| flgJ; peptidoglycan hydrolase FlgJ | K02395 | HC | 2.89 | 0.00 |
| flgI; flagellar P-ring protein FlgI | K02394 | HC | 2.44 | 0.00 |
| flgL; flagellar hook-associated protein 3 FlgL | K02397 | HC | 3.23 | 0.02 |
| flgK; flagellar hook-associated protein 1 | K02396 | HC | 3.45 | 0.04 |
| flgF; flagellar basal-body rod protein FlgF | K02391 | HC | 2.30 | 0.01 |
| flgE; flagellar hook protein FlgE | K02390 | HC | 3.43 | 0.02 |
| flgH; flagellar L-ring protein FlgH | K02393 | HC | 2.43 | 0.00 |
| flgG; flagellar basal-body rod protein FlgG | K02392 | HC | 3.63 | 0.04 |
| stpA; DNA-binding protein StpA | K11685 | AxSpA | 2.39 | 0.02 |
| catB; chloramphenicol O-acetyltransferase type B [EC:2.3.1.28] | K00638 | HC | 2.24 | 0.02 |
| mtnK; 5-methylthioribose kinase [EC:2.7.1.100] | K00899 | HC | 2.16 | 0.02 |
| gsk; inosine kinase [EC:2.7.1.73] | K00892 | AxSpA | 2.29 | 0.02 |
| int; integrase | K14059 | HC | 2.28 | 0.02 |
| E4.6.1.1; adenylate cyclase [EC:4.6.1.1] | K01768 | HC | 2.76 | 0.02 |
| RTCB, rtcB; tRNA-splicing ligase RtcB (3'-phosphate/5'-hydroxy nucleic acid ligase) [EC:6.5.1.8] | K14415 | HC | 2.68 | 0.02 |
| hoxH; NAD-reducing hydrogenase large subunit [EC:1.12.1.2] | K00436 | HC | 2.80 | 0.01 |
| K06884; uncharacterized protein | K06884 | AxSpA | 2.32 | 0.03 |
| cobQ, cbiP; adenosylcobyric acid synthase [EC:6.3.5.10] | K02232 | HC | 3.69 | 0.03 |
| E2.7.8.26, cobS, cobV; adenosylcobinamide-GDP ribazoletransferase [EC:2.7.8.26] | K02233 | HC | 3.67 | 0.02 |
| cobP, cobU; adenosylcobinamide kinase / adenosylcobinamide-phosphate guanylyltransferase [EC:2.7.1.156 2.7.7.62] | K02231 | HC | 3.77 | 0.03 |
| K16153; glycogen phosphorylase/synthase [EC:2.4.1.1 2.4.1.11] | K16153 | HC | 2.38 | 0.00 |
| hxpA; mannitol-1-/sugar-/sorbitol-6-phosphatase [EC:3.1.3.22 3.1.3.23 3.1.3.50] | K19270 | AxSpA | 2.19 | 0.01 |
| E1.2.7.8; indolepyruvate ferredoxin oxidoreductase [EC:1.2.7.8] | K04090 | HC | 2.07 | 0.03 |
| chrR, NQR; chromate reductase, NAD(P)H dehydrogenase (quinone) | K19784 | AxSpA | 3.63 | 0.05 |
| rof; Rho-binding antiterminator | K19000 | AxSpA | 2.23 | 0.04 |
| aroKB; shikimate kinase / 3-dehydroquinate synthase [EC:2.7.1.71 4.2.3.4] | K13829 | HC | 2.22 | 0.00 |
| CHAF1A; chromatin assembly factor 1 subunit A | K10750 | HC | 2.63 | 0.03 |
| PCCA, pccA; propionyl-CoA carboxylase alpha chain [EC:6.4.1.3] | K01965 | HC | 2.12 | 0.02 |
| legI, neuB2; N,N'-diacetyllegionaminate synthase [EC:2.5.1.101] | K18430 | HC | 3.00 | 0.05 |
| E6.4.1.4A; 3-methylcrotonyl-CoA carboxylase alpha subunit [EC:6.4.1.4] | K01968 | HC | 2.06 | 0.03 |
| parD1_3_4; antitoxin ParD1/3/4 | K07746 | HC | 2.36 | 0.02 |
| hutF; formimidoylglutamate deiminase [EC:3.5.3.13] | K05603 | HC | 2.07 | 0.02 |
| fadH; 2,4-dienoyl-CoA reductase (NADPH2) [EC:1.3.1.34] | K00219 | HC | 2.22 | 0.04 |
| coaX; type III pantothenate kinase [EC:2.7.1.33] | K03525 | HC | 3.73 | 0.02 |
| CRLS; cardiolipin synthase (CMP-forming) [EC:2.7.8.41] | K08744 | HC | 3.13 | 0.01 |
| acrF; multidrug efflux pump | K18142 | AxSpA | 2.28 | 0.02 |
| envR, acrS; TetR/AcrR family transcriptional regulator, acrEF/envCD operon repressor | K18140 | AxSpA | 2.28 | 0.02 |
| aroM; protein AroM | K14591 | AxSpA | 2.73 | 0.02 |
| rcnA; nickel/cobalt transporter (NicO) family protein | K08970 | AxSpA | 2.22 | 0.02 |
| wzxC; lipopolysaccharide exporter | K16695 | AxSpA | 2.60 | 0.05 |
| ecnB; entericidin B | K16348 | AxSpA | 2.57 | 0.05 |
| tqsA; AI-2 transport protein TqsA | K11744 | HC | 2.64 | 0.03 |
| AACS, acsA; acetoacetyl-CoA synthetase [EC:6.2.1.16] | K01907 | HC | 2.10 | 0.03 |
| ACSS3, prpE; propionyl-CoA synthetase [EC:6.2.1.17] | K01908 | HC | 2.08 | 0.02 |
| aroH; chorismate mutase [EC:5.4.99.5] | K06208 | HC | 2.32 | 0.01 |
| umuD; DNA polymerase V [EC:3.4.21.-] | K03503 | HC | 2.93 | 0.01 |
| cpaB, rcpC; pilus assembly protein CpaB | K02279 | HC | 2.66 | 0.01 |
| coxA, ctaD; cytochrome c oxidase subunit I [EC:7.1.1.9] | K02274 | HC | 2.58 | 0.03 |
| coxB, ctaC; cytochrome c oxidase subunit II [EC:7.1.1.9] | K02275 | HC | 2.58 | 0.03 |
| hisM; histidine transport system permease protein | K10015 | AxSpA | 2.40 | 0.02 |
| hisJ; histidine transport system substrate-binding protein | K10014 | AxSpA | 2.34 | 0.02 |
| hisP; histidine transport system ATP-binding protein [EC:7.4.2.1] | K10017 | AxSpA | 2.40 | 0.02 |
| hisQ; histidine transport system permease protein | K10016 | AxSpA | 2.40 | 0.02 |
| argT; lysine/arginine/ornithine transport system substrate-binding protein | K10013 | AxSpA | 2.03 | 0.04 |
| aac6-I, aacA7; aminoglycoside 6'-N-acetyltransferase I [EC:2.3.1.82] | K18816 | HC | 2.91 | 0.01 |
| K09925; uncharacterized protein | K09925 | HC | 2.33 | 0.02 |
| rcsA; LuxR family transcriptional regulator, capsular biosynthesis positive transcription factor | K07781 | AxSpA | 2.28 | 0.02 |
| sdiA; LuxR family transcriptional regulator, quorum-sensing system regulator SdiA | K07782 | AxSpA | 2.52 | 0.03 |
| G6PD, zwf; glucose-6-phosphate 1-dehydrogenase [EC:1.1.1.49 1.1.1.363] | K00036 | AxSpA | 3.75 | 0.04 |
| PGD, gnd, gntZ; 6-phosphogluconate dehydrogenase [EC:1.1.1.44 1.1.1.343] | K00033 | AxSpA | 3.77 | 0.02 |
| K09005; uncharacterized protein | K09005 | HC | 2.65 | 0.00 |
| IVD, ivd; isovaleryl-CoA dehydrogenase [EC:1.3.8.4] | K00253 | HC | 2.06 | 0.04 |
| ICP; inhibitor of cysteine peptidase | K14475 | HC | 2.06 | 0.02 |
| ftnB; ferritin-like protein 2 | K02255 | AxSpA | 2.28 | 0.02 |
| COX15, ctaA; heme a synthase [EC:1.17.99.9] | K02259 | HC | 2.57 | 0.05 |
| metL; bifunctional aspartokinase / homoserine dehydrogenase 2 [EC:2.7.2.4 1.1.1.3] | K12525 | AxSpA | 2.29 | 0.02 |
| eutR; AraC family transcriptional regulator, ethanolamine operon transcriptional activator | K04033 | AxSpA | 2.76 | 0.05 |
| nifD; nitrogenase molybdenum-iron protein alpha chain [EC:1.18.6.1] | K02586 | HC | 2.24 | 0.01 |
| phbB; acetoacetyl-CoA reductase [EC:1.1.1.36] | K00023 | HC | 2.21 | 0.03 |
| uspB; universal stress protein B | K06144 | AxSpA | 2.28 | 0.02 |
| tsgA; MFS transporter, TsgA protein | K06141 | AxSpA | 2.28 | 0.02 |
| csrD; RNase E specificity factor CsrD | K18765 | AxSpA | 2.24 | 0.02 |
| CHAC, chaC; glutathione-specific gamma-glutamylcyclotransferase [EC:4.3.2.7] | K07232 | HC | 2.30 | 0.02 |
| E1.1.1.30, bdh; 3-hydroxybutyrate dehydrogenase [EC:1.1.1.30] | K00019 | AxSpA | 2.69 | 0.03 |
| hpaA; AraC family transcriptional regulator, 4-hydroxyphenylacetate 3-monooxygenase operon regulatory protein | K02508 | AxSpA | 2.45 | 0.04 |
| rcsF; RcsF protein | K06080 | AxSpA | 2.24 | 0.02 |
| K09859; uncharacterized protein | K09859 | HC | 2.51 | 0.01 |
| dsdX; D-serine transporter | K13629 | AxSpA | 2.47 | 0.02 |
| acrE; membrane fusion protein, multidrug efflux system | K18141 | AxSpA | 2.31 | 0.02 |
| glsA, GLS; glutaminase [EC:3.5.1.2] | K01425 | HC | 2.94 | 0.04 |
| ghrA; glyoxylate/hydroxypyruvate reductase [EC:1.1.1.79 1.1.1.81] | K12972 | HC | 2.27 | 0.03 |
| pagP, crcA; lipid IVA palmitoyltransferase [EC:2.3.1.251] | K12973 | AxSpA | 2.48 | 0.04 |
| lpxP; KDO2-lipid IV(A) palmitoleoyltransferase [EC:2.3.1.242] | K12974 | AxSpA | 2.28 | 0.02 |
| hofM; pilus assembly protein HofM | K12288 | AxSpA | 2.43 | 0.02 |
| hofN; pilus assembly protein HofN | K12289 | AxSpA | 2.28 | 0.02 |
| urtA; urea transport system substrate-binding protein | K11959 | HC | 2.42 | 0.01 |
| hr; hemerythrin | K07216 | HC | 3.37 | 0.03 |
| hdhA; 7-alpha-hydroxysteroid dehydrogenase [EC:1.1.1.159] | K00076 | HC | 2.72 | 0.05 |
| paaH, hbd, fadB, mmgB; 3-hydroxybutyryl-CoA dehydrogenase [EC:1.1.1.157] | K00074 | HC | 3.34 | 0.02 |
| gfrA; fructoselysine/glucoselysine PTS system EIIA component [EC:2.7.1.-] | K19506 | HC | 2.69 | 0.01 |
| gfrB; fructoselysine/glucoselysine PTS system EIIB component [EC:2.7.1.-] | K19507 | HC | 2.71 | 0.02 |
| dexB; glucan 1,6-alpha-glucosidase [EC:3.2.1.70] | K01215 | AxSpA | 3.23 | 0.02 |
| gmuG; mannan endo-1,4-beta-mannosidase [EC:3.2.1.78] | K01218 | HC | 3.70 | 0.05 |
| mqnE; aminodeoxyfutalosine synthase [EC:2.5.1.120] | K18285 | HC | 2.41 | 0.00 |
| hycF; formate hydrogenlyase subunit 6 | K15831 | AxSpA | 2.47 | 0.02 |
| hycA; formate hydrogenlyase regulatory protein HycA | K15833 | AxSpA | 2.28 | 0.02 |
| mprA; two-component system, OmpR family, response regulator MprA | K07669 | HC | 2.43 | 0.02 |
| croR; 3-hydroxybutyryl-CoA dehydratase [EC:4.2.1.55] | K17865 | HC | 3.01 | 0.01 |
| pct; propionate CoA-transferase [EC:2.8.3.1] | K01026 | HC | 2.96 | 0.01 |
| hcr; NADH oxidoreductase Hcr [EC:1.-.-.-] | K11933 | AxSpA | 2.24 | 0.02 |
| uspG; universal stress protein G | K11932 | AxSpA | 2.10 | 0.03 |
| pgaA; biofilm PGA synthesis protein PgaA | K11935 | AxSpA | 2.56 | 0.05 |
| pgaD; biofilm PGA synthesis protein PgaD | K11937 | AxSpA | 2.56 | 0.04 |
| cof; HMP-PP phosphatase [EC:3.6.1.-] | K11938 | AxSpA | 2.28 | 0.02 |
| rssB, hnr; two-component system, response regulator | K02485 | AxSpA | 2.39 | 0.03 |
| fryB; fructose-like PTS system EIIB component [EC:2.7.1.-] | K11202 | HC | 2.78 | 0.01 |
| fruB; fructose PTS system EIIA component [EC:2.7.1.202] | K02768 | HC | 3.05 | 0.03 |
| fruAb; fructose PTS system EIIB component [EC:2.7.1.202] | K02769 | HC | 2.89 | 0.02 |
| gamP; D-glucosamine PTS system EIICBA component [EC:2.7.1.-] | K02765 | HC | 3.32 | 0.01 |
| licR; lichenan operon transcriptional antiterminator | K03491 | HC | 3.11 | 0.03 |
| basS; two-component system, OmpR family, sensor histidine kinase BasS [EC:2.7.13.3] | K07643 | AxSpA | 2.28 | 0.02 |
| E3.2.1.4; endoglucanase [EC:3.2.1.4] | K01179 | HC | 3.81 | 0.05 |
| phbC, phaC; polyhydroxyalkanoate synthase subunit PhaC [EC:2.3.1.-] | K03821 | HC | 2.42 | 0.02 |
| patA, rscA, lmrC, satA; ATP-binding cassette, subfamily B, multidrug efflux pump | K18891 | AxSpA | 3.14 | 0.04 |
| mdlB, smdB; ATP-binding cassette, subfamily B, multidrug efflux pump | K18890 | AxSpA | 2.24 | 0.02 |
| patB, rscB, lmrC, satB; ATP-binding cassette, subfamily B, multidrug efflux pump | K18892 | AxSpA | 3.13 | 0.04 |
| thiK; thiamine kinase [EC:2.7.1.89] | K07251 | AxSpA | 2.24 | 0.02 |
| spsF; spore coat polysaccharide biosynthesis protein SpsF | K07257 | HC | 2.22 | 0.04 |
| fliH; flagellar assembly protein FliH | K02411 | HC | 3.02 | 0.02 |
| mexK; multidrug efflux pump | K18303 | HC | 2.33 | 0.02 |
| ureJ; urease accessory protein | K03192 | HC | 2.15 | 0.01 |
| lmrS; MFS transporter, DHA2 family, multidrug resistance protein | K18934 | AxSpA | 2.41 | 0.03 |
| K09701; uncharacterized protein | K09701 | HC | 2.13 | 0.01 |
| meh; 3-methylfumaryl-CoA hydratase [EC:4.2.1.153] | K09709 | HC | 2.08 | 0.04 |
| bssR; biofilm regulator BssR | K19688 | AxSpA | 2.28 | 0.02 |
| cyaB; adenylate cyclase, class 2 [EC:4.6.1.1] | K05873 | AxSpA | 3.08 | 0.02 |
| cheR; chemotaxis protein methyltransferase CheR [EC:2.1.1.80] | K00575 | HC | 3.32 | 0.05 |
| jag; spoIIIJ-associated protein | K06346 | HC | 3.45 | 0.03 |
| norV; anaerobic nitric oxide reductase flavorubredoxin | K12264 | AxSpA | 2.24 | 0.02 |
| norW; nitric oxide reductase FlRd-NAD(+) reductase [EC:1.18.1.-] | K12265 | AxSpA | 2.24 | 0.02 |
| hycB; formate hydrogenlyase subunit 2 | K15827 | AxSpA | 2.28 | 0.02 |
| hycD; formate hydrogenlyase subunit 4 | K15829 | AxSpA | 2.28 | 0.02 |
| hycC; formate hydrogenlyase subunit 3 | K15828 | AxSpA | 2.28 | 0.02 |
| K09919; uncharacterized protein | K09919 | HC | 2.42 | 0.02 |
| ppnP; purine/pyrimidine-nucleoside phosphorylase [EC:2.4.2.1 2.4.2.2] | K09913 | HC | 2.35 | 0.01 |
| eamB; cysteine/O-acetylserine efflux protein | K11249 | HC | 2.42 | 0.00 |
| mdtD; MFS transporter, DHA2 family, multidrug resistance protein | K18326 | AxSpA | 2.28 | 0.02 |
| ramA; AraC family of transcriptional regulator, multidrug resistance transcriptional activator | K18325 | AxSpA | 2.43 | 0.04 |
| acrD; multidrug efflux pump | K18324 | AxSpA | 2.28 | 0.02 |
| IS15, IS26; transposase, IS6 family | K18320 | HC | 2.74 | 0.04 |
| aaeA; p-hydroxybenzoic acid efflux pump subunit AaeA | K15548 | AxSpA | 2.28 | 0.02 |
| trpGD; anthranilate synthase/phosphoribosyltransferase [EC:4.1.3.27 2.4.2.18] | K13497 | HC | 2.72 | 0.01 |
| fdnH; formate dehydrogenase-N, beta subunit | K08349 | AxSpA | 2.28 | 0.02 |
| aceB, glcB; malate synthase [EC:2.3.3.9] | K01638 | HC | 2.26 | 0.02 |
| nifH; nitrogenase iron protein NifH | K02588 | HC | 2.82 | 0.01 |
| puuP; putrescine importer | K14052 | AxSpA | 2.32 | 0.02 |
| rhaR; AraC family transcriptional regulator, L-rhamnose operon transcriptional activator RhaR | K02854 | AxSpA | 2.43 | 0.04 |
| whiA; cell division protein WhiA | K09762 | HC | 3.43 | 0.05 |
| cpxP, spy; periplasmic protein CpxP/Spy | K06006 | AxSpA | 2.31 | 0.02 |
| ymdB; 2',3'-cyclic-nucleotide 2'-phosphodiesterase [EC:3.1.4.16] | K09769 | HC | 3.11 | 0.02 |
| fimI; fimbrial protein | K07351 | AxSpA | 2.22 | 0.03 |
| K06915; uncharacterized protein | K06915 | AxSpA | 3.74 | 0.04 |
| K00375; GntR family transcriptional regulator / MocR family aminotransferase | K00375 | HC | 3.59 | 0.04 |
| phaZ; poly(3-hydroxybutyrate) depolymerase [EC:3.1.1.75] | K05973 | HC | 2.32 | 0.01 |
| ATPVA, ntpA, atpA; V/A-type H+/Na+-transporting ATPase subunit A [EC:7.1.2.2 7.2.2.1] | K02117 | HC | 3.72 | 0.03 |
| ATPVC, ntpC, atpC; V/A-type H+/Na+-transporting ATPase subunit C | K02119 | HC | 3.52 | 0.02 |
| ATPVB, ntpB, atpB; V/A-type H+/Na+-transporting ATPase subunit B | K02118 | HC | 3.75 | 0.03 |
| pgpC; phosphatidylglycerophosphatase C [EC:3.1.3.27] | K18697 | AxSpA | 2.46 | 0.02 |
| K07161; uncharacterized protein | K07161 | HC | 2.10 | 0.03 |
| nudG; (d)CTP diphosphatase [EC:3.6.1.65] | K08320 | AxSpA | 2.28 | 0.02 |
| K06973; uncharacterized protein | K06973 | HC | 3.61 | 0.05 |
| ATE1; arginyl-tRNA---protein transferase [EC:2.3.2.8] | K00685 | HC | 2.40 | 0.00 |
| hutC; GntR family transcriptional regulator, histidine utilization repressor | K05836 | HC | 2.11 | 0.02 |
| hha; haemolysin expression modulating protein | K05839 | AxSpA | 2.22 | 0.02 |
| alaA; alanine-synthesizing transaminase [EC:2.6.1.66 2.6.1.2] | K14260 | AxSpA | 3.72 | 0.04 |
| puuE; 4-aminobutyrate aminotransferase [EC:2.6.1.19] | K00823 | HC | 2.09 | 0.03 |
| E2.6.1.18; beta-alanine--pyruvate transaminase [EC:2.6.1.18] | K00822 | HC | 2.36 | 0.00 |
| add, ADA; adenosine deaminase [EC:3.5.4.4] | K01488 | HC | 3.10 | 0.01 |
| gutQ; arabinose 5-phosphate isomerase [EC:5.3.1.13] | K02467 | AxSpA | 2.28 | 0.02 |
| mtnD, mtnZ, ADI1; 1,2-dihydroxy-3-keto-5-methylthiopentene dioxygenase [EC:1.13.11.53 1.13.11.54] | K08967 | AxSpA | 2.18 | 0.04 |
| fliE; flagellar hook-basal body complex protein FliE | K02408 | HC | 3.35 | 0.04 |
| frsA; esterase FrsA [EC:3.1.-.-] | K11750 | AxSpA | 2.24 | 0.02 |
| gfrC; fructoselysine/glucoselysine PTS system EIIC component | K19508 | HC | 2.77 | 0.02 |
| gfrD; fructoselysine/glucoselysine PTS system EIID component | K19509 | HC | 2.70 | 0.02 |
| fliD; flagellar hook-associated protein 2 | K02407 | HC | 3.36 | 0.03 |
| fliC, hag; flagellin | K02406 | HC | 3.64 | 0.03 |
| E2.1.3.1-12S; methylmalonyl-CoA carboxyltransferase 12S subunit [EC:2.1.3.1] | K17489 | HC | 2.39 | 0.02 |
| iap; alkaline phosphatase isozyme conversion protein [EC:3.4.11.-] | K09612 | AxSpA | 2.52 | 0.02 |
| odh; opine dehydrogenase [EC:1.5.1.28] | K04940 | HC | 2.06 | 0.05 |
| msrP; methionine sulfoxide reductase catalytic subunit [EC:1.8.-.-] | K07147 | AxSpA | 2.69 | 0.02 |
| K07140; uncharacterized protein | K07140 | AxSpA | 2.69 | 0.04 |
| gatY-kbaY; tagatose 1,6-diphosphate aldolase GatY/KbaY [EC:4.1.2.40] | K08302 | HC | 3.03 | 0.02 |
| mltE, emtA; membrane-bound lytic murein transglycosylase E [EC:4.2.2.-] | K08308 | AxSpA | 2.28 | 0.02 |
| cynT, can; carbonic anhydrase [EC:4.2.1.1] | K01673 | AxSpA | 4.00 | 0.04 |
| E4.2.1.2AA, fumA; fumarate hydratase subunit alpha [EC:4.2.1.2] | K01677 | HC | 3.59 | 0.05 |
| E4.2.1.2AB, fumB; fumarate hydratase subunit beta [EC:4.2.1.2] | K01678 | HC | 3.59 | 0.05 |
| cheD; chemotaxis protein CheD [EC:3.5.1.44] | K03411 | HC | 3.29 | 0.04 |
| cheY; two-component system, chemotaxis family, chemotaxis protein CheY | K03413 | HC | 3.55 | 0.02 |
| cheB; two-component system, chemotaxis family, protein-glutamate methylesterase/glutaminase [EC:3.1.1.61 3.5.1.44] | K03412 | HC | 3.17 | 0.03 |
| cheZ; chemotaxis protein CheZ | K03414 | HC | 2.29 | 0.02 |
| msyB; acidic protein MsyB | K12147 | AxSpA | 2.28 | 0.02 |
| dinI; DNA-damage-inducible protein I | K12149 | AxSpA | 2.24 | 0.02 |
| bssS; biofilm regulator BssS | K12148 | AxSpA | 2.28 | 0.02 |
| K07317; adenine-specific DNA-methyltransferase [EC:2.1.1.72] | K07317 | HC | 2.41 | 0.01 |
| virB6, lvhB6; type IV secretion system protein VirB6 | K03201 | AxSpA | 2.36 | 0.04 |
| E2.7.13.3; histidine kinase [EC:2.7.13.3] | K10819 | AxSpA | 3.21 | 0.04 |
| K09124; uncharacterized protein | K09124 | HC | 2.50 | 0.02 |
| fliF; flagellar M-ring protein FliF | K02409 | HC | 3.25 | 0.03 |
| fre, ubiB; NAD(P)H-flavin reductase [EC:1.5.1.41] | K05368 | AxSpA | 2.29 | 0.02 |
| flhB; flagellar biosynthesis protein FlhB | K02401 | HC | 3.35 | 0.02 |
| flhA; flagellar biosynthesis protein FlhA | K02400 | HC | 3.35 | 0.05 |
| fliA, whiG; RNA polymerase sigma factor FliA | K02405 | HC | 3.34 | 0.05 |
| flhF; flagellar biosynthesis protein FlhF | K02404 | HC | 3.01 | 0.03 |
| PM20D1; carboxypeptidase PM20D1 [EC:3.4.17.-] | K13049 | HC | 3.14 | 0.04 |
| E1.14.13.40; anthraniloyl-CoA monooxygenase [EC:1.14.13.40] | K09461 | AxSpA | 2.55 | 0.01 |
| RYR2; ryanodine receptor 2 | K04962 | HC | 2.82 | 0.00 |
| K07129; uncharacterized protein | K07129 | AxSpA | 2.63 | 0.05 |
| ydgD; protease YdgD [EC:3.4.21.-] | K04775 | AxSpA | 2.28 | 0.02 |
| paaF, echA; enoyl-CoA hydratase [EC:4.2.1.17] | K01692 | HC | 2.49 | 0.01 |
| holE; DNA polymerase III subunit theta [EC:2.7.7.7] | K02345 | AxSpA | 2.22 | 0.02 |
| tehB; tellurite methyltransferase [EC:2.1.1.265] | K16868 | AxSpA | 3.78 | 0.01 |
| K07336; PKHD-type hydroxylase [EC:1.14.11.-] | K07336 | HC | 2.14 | 0.01 |
| ABC-2.LPSE.A; lipopolysaccharide transport system ATP-binding protein | K09691 | HC | 3.34 | 0.02 |
| ABC-2.LPSE.P; lipopolysaccharide transport system permease protein | K09690 | HC | 3.37 | 0.01 |
| tdcA; LysR family transcriptional regulator, tdc operon transcriptional activator | K07592 | AxSpA | 2.19 | 0.04 |
| sufA; Fe-S cluster assembly protein SufA | K05997 | AxSpA | 2.28 | 0.03 |
| ygiF; triphosphatase [EC:3.6.1.25] | K18446 | AxSpA | 2.28 | 0.01 |
| livM; branched-chain amino acid transport system permease protein | K01998 | HC | 3.68 | 0.02 |
| livK; branched-chain amino acid transport system substrate-binding protein | K01999 | HC | 3.69 | 0.01 |
| livF; branched-chain amino acid transport system ATP-binding protein | K01996 | HC | 3.65 | 0.01 |
| livH; branched-chain amino acid transport system permease protein | K01997 | HC | 3.63 | 0.04 |
| livG; branched-chain amino acid transport system ATP-binding protein | K01995 | HC | 3.66 | 0.02 |
| DAK, TKFC; triose/dihydroxyacetone kinase / FAD-AMP lyase (cyclizing) [EC:2.7.1.28 2.7.1.29 4.6.1.15] | K00863 | AxSpA | 3.13 | 0.03 |
| treS; maltose alpha-D-glucosyltransferase / alpha-amylase [EC:5.4.99.16 3.2.1.1] | K05343 | HC | 2.91 | 0.01 |
| K09726; uncharacterized protein | K09726 | HC | 2.41 | 0.05 |
| lnuA_C_D_E, lin; lincosamide nucleotidyltransferase A/C/D/E | K19545 | AxSpA | 2.46 | 0.01 |
| artJ; arginine transport system substrate-binding protein | K09996 | AxSpA | 2.00 | 0.03 |
| tupC, vupC; tungstate transport system ATP-binding protein [EC:7.3.2.6] | K06857 | HC | 2.34 | 0.01 |
| vsr; DNA mismatch endonuclease, patch repair protein [EC:3.1.-.-] | K07458 | HC | 3.27 | 0.01 |
| pcaB; 3-carboxy-cis,cis-muconate cycloisomerase [EC:5.5.1.2] | K01857 | HC | 2.22 | 0.01 |
| betA, CHDH; choline dehydrogenase [EC:1.1.99.1] | K00108 | HC | 2.28 | 0.03 |
| glcD; glycolate oxidase [EC:1.1.3.15] | K00104 | HC | 3.23 | 0.01 |
| LDHD, dld; D-lactate dehydrogenase (cytochrome) [EC:1.1.2.4] | K00102 | HC | 2.23 | 0.04 |
| sstT; serine/threonine transporter | K07862 | AxSpA | 3.74 | 0.04 |
| K00666; fatty-acyl-CoA synthase [EC:6.2.1.-] | K00666 | HC | 3.06 | 0.02 |
| tlyA; 23S rRNA (cytidine1920-2'-O)/16S rRNA (cytidine1409-2'-O)-methyltransferase [EC:2.1.1.226 2.1.1.227] | K06442 | HC | 3.47 | 0.04 |
| K07395; putative proteasome-type protease | K07395 | HC | 2.43 | 0.01 |
| ipdC; indolepyruvate decarboxylase [EC:4.1.1.74] | K04103 | AxSpA | 2.18 | 0.01 |
| glrR, qseF; two-component system, NtrC family, response regulator GlrR | K07715 | AxSpA | 2.26 | 0.01 |
| glrK, qseE; two-component system, NtrC family, sensor histidine kinase GlrK [EC:2.7.13.3] | K07711 | AxSpA | 2.31 | 0.01 |
| astC; succinylornithine aminotransferase [EC:2.6.1.81] | K00840 | AxSpA | 2.10 | 0.04 |
| HK; hexokinase [EC:2.7.1.1] | K00844 | HC | 2.51 | 0.01 |
| csiD; glutarate dioxygenase [EC:1.14.11.64] | K15737 | AxSpA | 2.32 | 0.02 |
| galP; MFS transporter, SP family, galactose:H+ symporter | K08137 | AxSpA | 2.28 | 0.02 |
| marR; MarR family transcriptional regulator, multiple antibiotic resistance protein MarR | K03712 | AxSpA | 2.28 | 0.02 |
| pheP; phenylalanine-specific permease | K11732 | AxSpA | 2.67 | 0.05 |
| proY; proline-specific permease ProY | K11736 | AxSpA | 2.28 | 0.02 |
| aroP; aromatic amino acid transport protein AroP | K11734 | HC | 2.32 | 0.00 |
| sbmC; DNA gyrase inhibitor | K07470 | AxSpA | 2.28 | 0.02 |
| cdh; CDP-diacylglycerol pyrophosphatase [EC:3.6.1.26] | K01521 | AxSpA | 2.45 | 0.01 |
| qrtT; energy-coupling factor transport system substrate-specific component | K16923 | HC | 3.33 | 0.02 |
| flgD; flagellar basal-body rod modification protein FlgD | K02389 | HC | 3.26 | 0.03 |
| flgB; flagellar basal-body rod protein FlgB | K02387 | HC | 3.33 | 0.05 |
| alsE; D-allulose-6-phosphate 3-epimerase [EC:5.1.3.-] | K17195 | HC | 2.28 | 0.04 |
| atzD; cyanuric acid amidohydrolase [EC:3.5.2.15] | K03383 | HC | 2.40 | 0.04 |
| tolC, bepC, cyaE, raxC, sapF, rsaF, hasF; outer membrane protein | K12340 | HC | 3.35 | 0.04 |
| flaG; flagellar protein FlaG | K06603 | HC | 2.91 | 0.03 |
| glcE; glycolate oxidase FAD binding subunit | K11472 | HC | 2.32 | 0.02 |
| glcF; glycolate oxidase iron-sulfur subunit | K11473 | HC | 2.30 | 0.05 |
| arnB, pmrH; UDP-4-amino-4-deoxy-L-arabinose-oxoglutarate aminotransferase [EC:2.6.1.87] | K07806 | AxSpA | 2.31 | 0.02 |
| paiB; transcriptional regulator | K07734 | HC | 2.27 | 0.01 |
| tlyC; putative hemolysin | K03699 | HC | 3.66 | 0.04 |
| ABCB1, CD243; ATP-binding cassette, subfamily B (MDR/TAP), member 1 [EC:7.6.2.2] | K05658 | AxSpA | 2.59 | 0.01 |
| sra; stationary-phase-induced ribosome-associated protein | K02972 | AxSpA | 2.28 | 0.02 |
| emrD; MFS transporter, DHA1 family, 2-module integral membrane pump EmrD | K08154 | AxSpA | 2.24 | 0.02 |
| araJ; MFS transporter, DHA1 family, arabinose polymer utilization protein | K08156 | HC | 2.55 | 0.03 |
| pfkB; 6-phosphofructokinase 2 [EC:2.7.1.11] | K16370 | HC | 2.24 | 0.01 |
| badH; 2-hydroxycyclohexanecarboxyl-CoA dehydrogenase [EC:1.1.1.-] | K07535 | HC | 2.43 | 0.04 |
| csxA; exo-1,4-beta-D-glucosaminidase [EC:3.2.1.165] | K15855 | HC | 2.16 | 0.03 |
| assT; arylsulfate sulfotransferase [EC:2.8.2.22] | K01023 | HC | 2.56 | 0.01 |
| E2.8.3.5A, scoA; 3-oxoacid CoA-transferase subunit A [EC:2.8.3.5] | K01028 | HC | 2.36 | 0.02 |
| rhaM; L-rhamnose mutarotase [EC:5.1.3.32] | K03534 | HC | 3.26 | 0.04 |
| cobC1, cobC; cobalamin biosynthesis protein CobC | K02225 | HC | 2.33 | 0.01 |
| cobB-cbiA; cobyrinic acid a,c-diamide synthase [EC:6.3.5.9 6.3.5.11] | K02224 | HC | 3.59 | 0.03 |
| cbiB, cobD; adenosylcobinamide-phosphate synthase [EC:6.3.1.10] | K02227 | HC | 3.68 | 0.03 |
| cobC, phpB; alpha-ribazole phosphatase [EC:3.1.3.73] | K02226 | HC | 3.83 | 0.03 |
| zapC; cell division protein ZapC | K18657 | AxSpA | 2.24 | 0.02 |
| K08884; serine/threonine protein kinase, bacterial [EC:2.7.11.1] | K08884 | HC | 3.80 | 0.04 |
| kapB; kinase-associated protein B | K06347 | AxSpA | 2.49 | 0.04 |
| ibpB; molecular chaperone IbpB | K04081 | AxSpA | 2.28 | 0.02 |
| secM; secretion monitor | K13301 | AxSpA | 2.28 | 0.02 |
| ulaE, sgaU, sgbU; L-ribulose-5-phosphate 3-epimerase [EC:5.1.3.22] | K03079 | HC | 2.52 | 0.01 |
| K08961; chondroitin-sulfate-ABC endolyase/exolyase [EC:4.2.2.20 4.2.2.21] | K08961 | HC | 2.66 | 0.02 |
| COQ7; 3-demethoxyubiquinol 3-hydroxylase [EC:1.14.99.60] | K06134 | HC | 2.33 | 0.01 |
| tus, tau; DNA replication terminus site-binding protein | K10748 | AxSpA | 2.24 | 0.02 |
| paaK; phenylacetate-CoA ligase [EC:6.2.1.30] | K01912 | HC | 3.67 | 0.03 |
| basR; two-component system, OmpR family, response regulator BasR | K07771 | AxSpA | 2.28 | 0.02 |
| fabV, ter; enoyl-[acyl-carrier protein] reductase / trans-2-enoyl-CoA reductase (NAD+) [EC:1.3.1.9 1.3.1.44] | K00209 | HC | 2.93 | 0.00 |
| exuT; MFS transporter, ACS family, hexuronate transporter | K08191 | HC | 3.16 | 0.04 |
| thrH; phosphoserine / homoserine phosphotransferase [EC:3.1.3.3 2.7.1.39] | K02203 | HC | 3.11 | 0.03 |
| tadB; tight adherence protein B | K12510 | HC | 3.38 | 0.03 |
| tadC; tight adherence protein C | K12511 | HC | 2.75 | 0.02 |
| nagK; fumarylpyruvate hydrolase [EC:3.7.1.20] | K16165 | HC | 2.26 | 0.03 |
| cdgJ; c-di-GMP phosphodiesterase [EC:3.1.4.52] | K07181 | HC | 2.92 | 0.01 |
| K09803; uncharacterized protein | K09803 | HC | 2.29 | 0.03 |
| K09805; uncharacterized protein | K09805 | HC | 2.89 | 0.01 |
| flhB2; flagellar biosynthesis protein | K04061 | HC | 3.24 | 0.03 |
| apgM; 2,3-bisphosphoglycerate-independent phosphoglycerate mutase [EC:5.4.2.12] | K15635 | HC | 3.52 | 0.05 |
| cueO; blue copper oxidase | K14588 | AxSpA | 2.50 | 0.02 |
| dgs, bgsA; 1,2-diacylglycerol-3-alpha-glucose alpha-1,2-glucosyltransferase [EC:2.4.1.208] | K13677 | HC | 2.97 | 0.00 |
| fliS; flagellar secretion chaperone FliS | K02422 | HC | 3.46 | 0.04 |
| fliR; flagellar biosynthesis protein FliR | K02421 | HC | 3.35 | 0.04 |
| wapR; alpha-1,3-rhamnosyltransferase [EC:2.4.1.-] | K12988 | HC | 2.45 | 0.00 |
| APOD; apolipoprotein D and lipocalin family protein | K03098 | HC | 2.25 | 0.01 |
| K07577; putative mRNA 3-end processing factor | K07577 | HC | 2.04 | 0.04 |
| tctB; putative tricarboxylic transport membrane protein | K07794 | HC | 2.16 | 0.03 |
| E3.1.1.45; carboxymethylenebutenolidase [EC:3.1.1.45] | K01061 | HC | 2.63 | 0.02 |
| aes; acetyl esterase [EC:3.1.1.-] | K01066 | HC | 2.35 | 0.02 |
| cbiX; sirohydrochlorin cobaltochelatase [EC:4.99.1.3] | K03795 | HC | 2.43 | 0.01 |
| pksJ; polyketide synthase PksJ | K13611 | HC | 2.49 | 0.00 |
| terD; tellurium resistance protein TerD | K05795 | HC | 2.61 | 0.01 |
| hydN; electron transport protein HydN | K05796 | HC | 2.04 | 0.01 |
| leuO; LysR family transcriptional regulator, transcriptional activator for leuABCD operon | K05798 | AxSpA | 2.28 | 0.02 |
| TC.CITMHS; citrate-Mg2+:H+ or citrate-Ca2+:H+ symporter, CitMHS family | K03300 | HC | 2.46 | 0.03 |
| alkA; DNA-3-methyladenine glycosylase II [EC:3.2.2.21] | K01247 | HC | 2.36 | 0.05 |
| epsJ; glycosyltransferase EpsJ [EC:2.4.-.-] | K19427 | AxSpA | 2.29 | 0.04 |
| urtD; urea transport system ATP-binding protein | K11962 | HC | 2.41 | 0.01 |
| urtB; urea transport system permease protein | K11960 | HC | 2.42 | 0.01 |
| urtC; urea transport system permease protein | K11961 | HC | 2.42 | 0.01 |
| mutM, fpg; formamidopyrimidine-DNA glycosylase [EC:3.2.2.23 4.2.99.18] | K10563 | AxSpA | 3.75 | 0.04 |
| yqjH; ferric-chelate reductase (NADPH) [EC:1.16.1.9] | K07229 | AxSpA | 2.28 | 0.02 |
| frlC; fructoselysine 3-epimerase [EC:5.1.3.41] | K10709 | HC | 2.78 | 0.01 |
| rutR; TetR/AcrR family transcriptional regulator | K09017 | HC | 2.17 | 0.01 |
| DNPEP; aspartyl aminopeptidase [EC:3.4.11.21] | K01267 | HC | 3.52 | 0.02 |
| lplT; MFS transporter, LPLT family, lysophospholipid transporter | K08227 | AxSpA | 2.28 | 0.02 |
| entS; MFS transporter, ENTS family, enterobactin (siderophore) exporter | K08225 | AxSpA | 2.44 | 0.04 |
| tomB; hha toxicity modulator TomB | K19162 | AxSpA | 2.28 | 0.02 |
| comFB; competence protein ComFB | K02241 | HC | 2.80 | 0.03 |
| tauD; taurine dioxygenase [EC:1.14.11.17] | K03119 | HC | 2.30 | 0.02 |
| rlmG; 23S rRNA (guanine1835-N2)-methyltransferase [EC:2.1.1.174] | K11391 | AxSpA | 2.29 | 0.02 |
| PTH2; peptidyl-tRNA hydrolase, PTH2 family [EC:3.1.1.29] | K04794 | HC | 2.94 | 0.02 |
| soxS; AraC family transcriptional regulator, mar-sox-rob regulon activator | K13631 | AxSpA | 2.22 | 0.02 |
| marA; AraC family transcriptional regulator, mar-sox-rob regulon activator | K13632 | AxSpA | 2.19 | 0.04 |
| ftrA; AraC family transcriptional regulator, transcriptional activator FtrA | K13633 | AxSpA | 2.43 | 0.05 |
| ABC.VB1X.P; putative thiamine transport system permease protein | K05778 | AxSpA | 2.47 | 0.05 |
| ABC.VB1X.A; putative thiamine transport system ATP-binding protein | K05779 | AxSpA | 2.47 | 0.05 |
| tupA, vupA; tungstate transport system substrate-binding protein | K05772 | HC | 2.33 | 0.01 |
| tupB, vupB; tungstate transport system permease protein | K05773 | HC | 2.33 | 0.01 |
| tsx; nucleoside-specific channel-forming protein | K05517 | AxSpA | 2.30 | 0.02 |
| nudK; GDP-mannose pyrophosphatase NudK [EC:3.6.1.-] | K12945 | AxSpA | 2.28 | 0.02 |
| ygeR; lipoprotein YgeR | K12943 | AxSpA | 2.36 | 0.02 |
| ACR3, arsB; arsenite transporter | K03325 | HC | 2.68 | 0.01 |
| mdoC; glucans biosynthesis protein C [EC:2.1.-.-] | K11941 | AxSpA | 2.28 | 0.02 |
| gldA; glycerol dehydrogenase [EC:1.1.1.6] | K00005 | HC | 3.27 | 0.03 |
| gltD; glutamate synthase (NADPH) small chain [EC:1.4.1.13] | K00266 | HC | 4.04 | 0.05 |
| gltB; glutamate synthase (NADPH) large chain [EC:1.4.1.13] | K00265 | HC | 3.18 | 0.03 |
| GBA, srfJ; glucosylceramidase [EC:3.2.1.45] | K01201 | HC | 2.84 | 0.02 |
| ynaI, mscMJ; MscS family membrane protein | K16052 | HC | 2.94 | 0.02 |
| K07075; uncharacterized protein | K07075 | HC | 2.97 | 0.01 |
| arnF; undecaprenyl phosphate-alpha-L-ara4N flippase subunit ArnF | K12963 | AxSpA | 2.31 | 0.02 |
| diaA; DnaA initiator-associating protein | K12961 | AxSpA | 2.29 | 0.02 |
| rzpD; prophage endopeptidase [EC:3.4.-.-] | K14744 | AxSpA | 2.42 | 0.02 |
| hndA; NADP-reducing hydrogenase subunit HndA [EC:1.12.1.3] | K18330 | HC | 3.45 | 0.05 |
| hndC; NADP-reducing hydrogenase subunit HndC [EC:1.12.1.3] | K18331 | HC | 3.44 | 0.04 |
| K18333; L-fucose dehydrogenase | K18333 | HC | 2.36 | 0.03 |
| pabBC; para-aminobenzoate synthetase / 4-amino-4-deoxychorismate lyase [EC:2.6.1.85 4.1.3.38] | K03342 | AxSpA | 3.66 | 0.02 |
| mngR, farR; GntR family transcriptional regulator, mannosyl-D-glycerate transport/metabolism system repressor | K11922 | HC | 2.13 | 0.03 |
| crl; sigma factor-binding protein Crl | K11926 | AxSpA | 2.24 | 0.02 |
| sgrR; SgrR family transcriptional regulator | K11925 | AxSpA | 2.38 | 0.02 |
| phoE; outer membrane pore protein E | K11929 | AxSpA | 2.28 | 0.02 |
| arnT, pmrK; 4-amino-4-deoxy-L-arabinose transferase [EC:2.4.2.43] | K07264 | AxSpA | 2.32 | 0.02 |
| ytfB; uncharacterized protein | K07269 | AxSpA | 2.24 | 0.02 |
| SHPK; sedoheptulokinase [EC:2.7.1.14] | K11214 | HC | 2.21 | 0.03 |
| clfB; clumping factor B | K14192 | AxSpA | 2.30 | 0.03 |
| E3.2.1.89; arabinogalactan endo-1,4-beta-galactosidase [EC:3.2.1.89] | K01224 | HC | 3.24 | 0.04 |
| mepH; murein DD-endopeptidase [EC:3.4.-.-] | K19303 | AxSpA | 2.28 | 0.02 |
| tdcD; propionate kinase [EC:2.7.2.15] | K00932 | AxSpA | 2.28 | 0.02 |
| treR; LacI family transcriptional regulator, trehalose operon repressor | K03485 | AxSpA | 2.15 | 0.04 |
| cpaE, tadZ; pilus assembly protein CpaE | K02282 | HC | 2.29 | 0.00 |
| cpaC, rcpA; pilus assembly protein CpaC | K02280 | HC | 2.28 | 0.02 |
| K17213; inositol transport system substrate-binding protein | K17213 | HC | 2.90 | 0.02 |
| rcsD; two-component system, NarL family, sensor histidine kinase RcsD [EC:2.7.13.3] | K07676 | AxSpA | 2.28 | 0.02 |
| xni; protein Xni | K01146 | AxSpA | 2.29 | 0.02 |
| yibL; ribosome-associated protein | K14762 | AxSpA | 2.28 | 0.02 |
| wcaB; putative colanic acid biosynthesis acetyltransferase WcaB [EC:2.3.1.-] | K03819 | AxSpA | 2.60 | 0.05 |
| aldA; lactaldehyde dehydrogenase / glycolaldehyde dehydrogenase [EC:1.2.1.22 1.2.1.21] | K07248 | AxSpA | 3.57 | 0.03 |
| ttuC, dmlA; tartrate dehydrogenase/decarboxylase / D-malate dehydrogenase [EC:1.1.1.93 4.1.1.73 1.1.1.83] | K07246 | HC | 2.53 | 0.01 |
| hoxN, nixA; nickel/cobalt transporter (NiCoT) family protein | K07241 | AxSpA | 2.66 | 0.02 |
| uxuB; fructuronate reductase [EC:1.1.1.57] | K00040 | HC | 3.13 | 0.04 |
| E1.1.1.67, mtlK; mannitol 2-dehydrogenase [EC:1.1.1.67] | K00045 | HC | 2.07 | 0.04 |
| nadM; nicotinamide-nucleotide adenylyltransferase [EC:2.7.7.1] | K00952 | HC | 2.14 | 0.01 |
| pcaC; 4-carboxymuconolactone decarboxylase [EC:4.1.1.44] | K01607 | HC | 3.68 | 0.01 |
| motA; chemotaxis protein MotA | K02556 | HC | 3.48 | 0.04 |
| wcaJ; putative colanic acid biosysnthesis UDP-glucose lipid carrier transferase | K03606 | HC | 3.15 | 0.05 |
| lpp; murein lipoprotein | K06078 | AxSpA | 2.25 | 0.03 |
| cutF, nlpE; copper homeostasis protein (lipoprotein) | K06079 | HC | 2.39 | 0.02 |
| btuC; vitamin B12 transport system permease protein | K06073 | AxSpA | 2.24 | 0.02 |
| btuD; vitamin B12 transport system ATP-binding protein [EC:7.6.2.8] | K06074 | AxSpA | 2.24 | 0.02 |
| ompR; two-component system, OmpR family, phosphate regulon response regulator OmpR | K07659 | HC | 2.47 | 0.03 |
| algA, xanB, rfbA, wbpW, pslB; mannose-1-phosphate guanylyltransferase / mannose-6-phosphate isomerase [EC:2.7.7.13 5.3.1.8] | K16011 | HC | 2.33 | 0.01 |
| K09153; small membrane protein | K09153 | HC | 3.12 | 0.03 |
| acpH; acyl carrier protein phosphodiesterase [EC:3.1.4.14] | K08682 | AxSpA | 2.28 | 0.02 |
| kdgT; 2-keto-3-deoxygluconate permease | K02526 | HC | 2.44 | 0.00 |
| ATPVF, ntpF, atpF; V/A-type H+/Na+-transporting ATPase subunit F | K02122 | HC | 3.65 | 0.01 |
| ATPVI, ntpI, atpI; V/A-type H+/Na+-transporting ATPase subunit I | K02123 | HC | 3.75 | 0.03 |
| ATPVD, ntpD, atpD; V/A-type H+/Na+-transporting ATPase subunit D | K02120 | HC | 3.75 | 0.03 |
| ATPVE, ntpE, atpE; V/A-type H+/Na+-transporting ATPase subunit E | K02121 | HC | 3.46 | 0.01 |
| ATPVK, ntpK, atpK; V/A-type H+/Na+-transporting ATPase subunit K | K02124 | HC | 3.75 | 0.03 |
| tdcC; threonine transporter | K03838 | AxSpA | 2.28 | 0.02 |
| fhuF; ferric iron reductase protein FhuF | K13255 | AxSpA | 2.22 | 0.05 |
| mqnC; cyclic dehypoxanthinyl futalosine synthase [EC:1.21.98.1] | K11784 | HC | 2.36 | 0.00 |
| mqnD; 1,4-dihydroxy-6-naphthoate synthase [EC:1.14.-.-] | K11785 | HC | 2.27 | 0.01 |
| cbe, mbe; cellobiose epimerase [EC:5.1.3.11] | K16213 | HC | 3.09 | 0.04 |
| mgp; 4-O-beta-D-mannosyl-D-glucose phosphorylase [EC:2.4.1.281] | K16212 | HC | 3.20 | 0.03 |
| ynhG; L,D-transpeptidase YnhG | K19234 | AxSpA | 2.28 | 0.02 |
| hndD; NADP-reducing hydrogenase subunit HndD [EC:1.12.1.3] | K18332 | HC | 3.46 | 0.04 |
| K18335; 2-keto-3-deoxy-L-fuconate dehydrogenase [EC:1.1.1.-] | K18335 | HC | 2.14 | 0.01 |
| E4.1.1.32, pckA, PCK; phosphoenolpyruvate carboxykinase (GTP) [EC:4.1.1.32] | K01596 | HC | 2.93 | 0.02 |
| ppc; phosphoenolpyruvate carboxylase [EC:4.1.1.31] | K01595 | AxSpA | 3.75 | 0.03 |
| hemE, UROD; uroporphyrinogen decarboxylase [EC:4.1.1.37] | K01599 | AxSpA | 3.76 | 0.01 |
| ltaE; threonine aldolase [EC:4.1.2.48] | K01620 | HC | 3.66 | 0.02 |
| rubB, alkT; rubredoxin---NAD+ reductase [EC:1.18.1.1] | K05297 | HC | 2.58 | 0.01 |
| mdoH; membrane glycosyltransferase [EC:2.4.1.-] | K03669 | HC | 2.10 | 0.05 |
| STE24; STE24 endopeptidase [EC:3.4.24.84] | K06013 | HC | 2.32 | 0.00 |
| envZ; two-component system, OmpR family, osmolarity sensor histidine kinase EnvZ [EC:2.7.13.3] | K07638 | HC | 2.31 | 0.01 |
| UMF1; MFS transporter, UMF1 family | K06902 | HC | 3.22 | 0.04 |

| F_ASvHC_IgA-_a0.05_w0.05_l2_minc.default.res.sig.txt | | | | |
| --- | --- | --- | --- | --- |
| Level D (KO)_Description | KO | Class with highest mean | Log LDA score | p-value (KW for class) |
| atoA; acetate CoA/acetoacetate CoA-transferase beta subunit [EC:2.8.3.8 2.8.3.9] | K01035 | HC | 2.66 | 0.03 |
| atoD; acetate CoA/acetoacetate CoA-transferase alpha subunit [EC:2.8.3.8 2.8.3.9] | K01034 | HC | 2.60 | 0.04 |
| lysY; putative lysine transport system ATP-binding protein | K17076 | HC | 2.52 | 0.05 |
| lysX2; putative lysine transport system permease protein | K17074 | HC | 2.52 | 0.04 |
| E3.5.1.4, amiE; amidase [EC:3.5.1.4] | K01426 | HC | 2.71 | 0.04 |
| wbpI, wlbD; UDP-GlcNAc3NAcA epimerase [EC:5.1.3.23] | K13019 | HC | 2.08 | 0.05 |
| PPCS, COAB; phosphopantothenate---cysteine ligase (ATP) [EC:6.3.2.51] | K01922 | HC | 2.39 | 0.03 |
| pksJ; polyketide synthase PksJ | K13611 | HC | 2.29 | 0.03 |
| abfD; 4-hydroxybutyryl-CoA dehydratase / vinylacetyl-CoA-Delta-isomerase [EC:4.2.1.120 5.3.3.3] | K14534 | HC | 2.58 | 0.03 |
| yahK; alcohol dehydrogenase (NADP+) [EC:1.1.1.2] | K13979 | HC | 2.09 | 0.03 |
| NEU1; sialidase-1 [EC:3.2.1.18] | K01186 | HC | 3.24 | 0.04 |
| cynR; LysR family transcriptional regulator, cyn operon transcriptional activator | K11921 | HC | 2.95 | 0.03 |
| gcvPB; glycine dehydrogenase subunit 2 [EC:1.4.4.2] | K00283 | HC | 2.56 | 0.03 |
| gcvPA; glycine dehydrogenase subunit 1 [EC:1.4.4.2] | K00282 | HC | 2.56 | 0.02 |
| gerKB; spore germination protein KB | K06296 | HC | 2.37 | 0.00 |
| gerKC; spore germination protein KC | K06297 | HC | 2.38 | 0.03 |
| PC, pyc; pyruvate carboxylase [EC:6.4.1.1] | K01958 | HC | 2.76 | 0.02 |
| ctsR; transcriptional regulator of stress and heat shock response | K03708 | HC | 2.95 | 0.03 |
| E1.11.1.5; cytochrome c peroxidase [EC:1.11.1.5] | K00428 | HC | 3.08 | 0.04 |
| mmsA, iolA, ALDH6A1; malonate-semialdehyde dehydrogenase (acetylating) / methylmalonate-semialdehyde dehydrogenase [EC:1.2.1.18 1.2.1.27] | K00140 | HC | 2.38 | 0.03 |
| iolC; 5-dehydro-2-deoxygluconokinase [EC:2.7.1.92] | K03338 | HC | 2.56 | 0.04 |
| epsJ; glycosyltransferase EpsJ [EC:2.4.-.-] | K19427 | HC | 2.18 | 0.04 |
| sspD; small acid-soluble spore protein D (minor alpha/beta-type SASP) | K06421 | HC | 2.24 | 0.03 |
| ppaX; pyrophosphatase PpaX [EC:3.6.1.1] | K06019 | HC | 2.61 | 0.04 |

| F_ASvHC_IgA+_a0.05_w0.05_l2_minc.default.res.sig.txt | | | | |
| --- | --- | --- | --- | --- |
| Level D (KO) _Description | KO | Class with highest mean | Log LDA score | p-value (KW for class) |
| ltrA; RNA-directed DNA polymerase [EC:2.7.7.49] | K00986 | AxSpA | 2.88 | 0.04 |
| pseB, wbjB; UDP-N-acetylglucosamine 4,6-dehydratase/5-epimerase [EC:4.2.1.115 5.1.3.-] | K15894 | AxSpA | 2.21 | 0.02 |
| TC.KEF; monovalent cation:H+ antiporter-2, CPA2 family | K03455 | AxSpA | 3.03 | 0.04 |
| ftnA, ftn; ferritin [EC:1.16.3.2] | K02217 | AxSpA | 3.21 | 0.05 |
| araN; arabinosaccharide transport system substrate-binding protein | K17234 | AxSpA | 2.43 | 0.01 |
| gmhA, lpcA; D-sedoheptulose 7-phosphate isomerase [EC:5.3.1.28] | K03271 | AxSpA | 3.01 | 0.03 |
| HYDIN; hydrocephalus-inducing protein | K17570 | AxSpA | 2.61 | 0.00 |
| asnA; aspartate--ammonia ligase [EC:6.3.1.1] | K01914 | AxSpA | 3.44 | 0.05 |
| cld; chlorite dismutase [EC:1.13.11.49] | K09162 | AxSpA | 2.61 | 0.01 |
| lonB; ATP-dependent Lon protease [EC:3.4.21.53] | K04076 | AxSpA | 2.62 | 0.00 |
| HMOX1; heme oxygenase 1 [EC:1.14.14.18] | K00510 | AxSpA | 2.62 | 0.00 |
| SMARCAL1, HARP; SWI/SNF-related matrix-associated actin-dependent regulator of chromatin subfamily A-like protein 1 [EC:3.6.4.12] | K14440 | AxSpA | 2.47 | 0.03 |
| SLC13A2_3_5; solute carrier family 13 (sodium-dependent dicarboxylate transporter), member 2/3/5 | K14445 | AxSpA | 2.86 | 0.00 |
| abnA; arabinan endo-1,5-alpha-L-arabinosidase [EC:3.2.1.99] | K06113 | AxSpA | 3.29 | 0.01 |
| aspA; aspartate ammonia-lyase [EC:4.3.1.1] | K01744 | AxSpA | 3.20 | 0.03 |
| assT; arylsulfate sulfotransferase [EC:2.8.2.22] | K01023 | AxSpA | 2.22 | 0.03 |
| tolA; colicin import membrane protein | K03646 | AxSpA | 2.83 | 0.02 |
| PPOX, hemY; protoporphyrinogen/coproporphyrinogen III oxidase [EC:1.3.3.4 1.3.3.15] | K00231 | AxSpA | 2.68 | 0.02 |
| sdhA, frdA; succinate dehydrogenase / fumarate reductase, flavoprotein subunit [EC:1.3.5.1 1.3.5.4] | K00239 | AxSpA | 3.09 | 0.04 |
| K02475; two-component system, CitB family, response regulator | K02475 | AxSpA | 2.43 | 0.01 |
| K02476; two-component system, CitB family, sensor kinase [EC:2.7.13.3] | K02476 | AxSpA | 2.51 | 0.02 |
| K07085; putative transport protein | K07085 | AxSpA | 3.44 | 0.05 |
| hipA; serine/threonine-protein kinase HipA [EC:2.7.11.1] | K07154 | AxSpA | 3.48 | 0.02 |
| araA; L-arabinose isomerase [EC:5.3.1.4] | K01804 | AxSpA | 3.27 | 0.04 |
| tag; DNA-3-methyladenine glycosylase I [EC:3.2.2.20] | K01246 | AxSpA | 3.12 | 0.02 |
| wecB; UDP-N-acetylglucosamine 2-epimerase (non-hydrolysing) [EC:5.1.3.14] | K01791 | AxSpA | 3.56 | 0.01 |
| htpX; heat shock protein HtpX [EC:3.4.24.-] | K03799 | AxSpA | 2.92 | 0.04 |
| cphB; cyanophycinase [EC:3.4.15.6] | K13282 | AxSpA | 2.61 | 0.00 |
| yfkQ; spore germination protein | K06307 | AxSpA | 2.61 | 0.00 |
| spoVR; stage V sporulation protein R | K06415 | AxSpA | 2.62 | 0.00 |
| K09133; uncharacterized protein | K09133 | AxSpA | 2.54 | 0.04 |
| xylS, yicI; alpha-D-xyloside xylohydrolase [EC:3.2.1.177] | K01811 | AxSpA | 3.51 | 0.02 |
| dps; starvation-inducible DNA-binding protein | K04047 | AxSpA | 2.97 | 0.04 |
| lip, TGL2; triacylglycerol lipase [EC:3.1.1.3] | K01046 | AxSpA | 2.37 | 0.04 |
| G6PD, zwf; glucose-6-phosphate 1-dehydrogenase [EC:1.1.1.49 1.1.1.363] | K00036 | AxSpA | 2.97 | 0.04 |
| PGD, gnd, gntZ; 6-phosphogluconate dehydrogenase [EC:1.1.1.44 1.1.1.343] | K00033 | AxSpA | 2.97 | 0.04 |
| manA, MPI; mannose-6-phosphate isomerase [EC:5.3.1.8] | K01809 | AxSpA | 3.27 | 0.02 |
| phaF; multicomponent K+:H+ antiporter subunit F | K05563 | AxSpA | 2.62 | 0.00 |
| nrfH; cytochrome c nitrite reductase small subunit | K15876 | AxSpA | 2.72 | 0.01 |
| rocD, OAT; ornithine--oxo-acid transaminase [EC:2.6.1.13] | K00819 | AxSpA | 2.24 | 0.02 |
| terD; tellurium resistance protein TerD | K05795 | AxSpA | 2.73 | 0.02 |
| E4.2.1.2B, fumC, FH; fumarate hydratase, class II [EC:4.2.1.2] | K01679 | AxSpA | 2.76 | 0.02 |
| rluA; tRNA pseudouridine32 synthase / 23S rRNA pseudouridine746 synthase [EC:5.4.99.28 5.4.99.29] | K06177 | AxSpA | 3.02 | 0.05 |
| lemA; LemA protein | K03744 | AxSpA | 3.19 | 0.03 |
| K07133; uncharacterized protein | K07133 | AxSpA | 3.97 | 0.05 |
| ihfA, himA; integration host factor subunit alpha | K04764 | AxSpA | 2.75 | 0.03 |
| STAR2, fetB; UDP-glucose/iron transport system permease protein | K02069 | AxSpA | 2.98 | 0.04 |
| STAR1, fetA; UDP-glucose/iron transport system ATP-binding protein | K02068 | AxSpA | 2.91 | 0.04 |
| phaG; multicomponent K+:H+ antiporter subunit G | K05564 | AxSpA | 2.62 | 0.00 |
| aguA; alpha-glucuronidase [EC:3.2.1.139] | K01235 | AxSpA | 2.85 | 0.02 |
| dptF; DNA phosphorothioation-dependent restriction protein DptF | K19173 | AxSpA | 2.35 | 0.04 |
| iadA; beta-aspartyl-dipeptidase (metallo-type) [EC:3.4.19.-] | K01305 | AxSpA | 2.71 | 0.05 |
| pcp; pyroglutamyl-peptidase [EC:3.4.19.3] | K01304 | AxSpA | 2.47 | 0.02 |
| PGLS, pgl, devB; 6-phosphogluconolactonase [EC:3.1.1.31] | K01057 | AxSpA | 2.81 | 0.05 |
| fabB; 3-oxoacyl-[acyl-carrier-protein] synthase I [EC:2.3.1.41] | K00647 | AxSpA | 3.08 | 0.01 |
| fitB; toxin FitB [EC:3.1.-.-] | K07062 | AxSpA | 2.58 | 0.04 |
| cysE; serine O-acetyltransferase [EC:2.3.1.30] | K00640 | AxSpA | 3.58 | 0.04 |
| ogl; oligogalacturonide lyase [EC:4.2.2.6] | K01730 | AxSpA | 2.25 | 0.02 |
| fklB; FKBP-type peptidyl-prolyl cis-trans isomerase FklB [EC:5.2.1.8] | K03773 | AxSpA | 3.45 | 0.03 |
| K07217; Mn-containing catalase | K07217 | AxSpA | 2.34 | 0.04 |
| ATOX1, ATX1, copZ, golB; copper chaperone | K07213 | AxSpA | 2.66 | 0.01 |
| mcrB; 5-methylcytosine-specific restriction enzyme B [EC:3.1.21.-] | K07452 | AxSpA | 2.54 | 0.04 |
| vanRB, vanR, vanRD; two-component system, OmpR family, response regulator VanR | K18344 | AxSpA | 2.61 | 0.04 |
| E3.2.1.8, xynA; endo-1,4-beta-xylanase [EC:3.2.1.8] | K01181 | AxSpA | 2.80 | 0.00 |
| ftsI; cell division protein FtsI (penicillin-binding protein 3) [EC:3.4.16.4] | K03587 | AxSpA | 3.12 | 0.04 |
| coxM, cutM; aerobic carbon-monoxide dehydrogenase medium subunit [EC:1.2.5.3] | K03519 | AxSpA | 2.23 | 0.04 |
| LIG1; DNA ligase 1 [EC:6.5.1.1 6.5.1.6 6.5.1.7] | K10747 | AxSpA | 2.13 | 0.04 |
| fic; cell filamentation protein | K04095 | AxSpA | 3.18 | 0.03 |
| slo; thiol-activated cytolysin | K11031 | AxSpA | 2.40 | 0.01 |
| pqqL; zinc protease [EC:3.4.24.-] | K07263 | AxSpA | 3.08 | 0.03 |
| treR2, treR; GntR family transcriptional regulator, trehalose operon transcriptional repressor | K03486 | AxSpA | 2.53 | 0.03 |
| cphA; cyanophycin synthetase [EC:6.3.2.29 6.3.2.30] | K03802 | AxSpA | 2.62 | 0.00 |
| acm; lysozyme | K07273 | AxSpA | 3.29 | 0.03 |
| gpx, btuE, bsaA; glutathione peroxidase [EC:1.11.1.9] | K00432 | AxSpA | 3.21 | 0.00 |
| cpo; non-heme chloroperoxidase [EC:1.11.1.10] | K00433 | AxSpA | 2.39 | 0.03 |
| pgpA; phosphatidylglycerophosphatase A [EC:3.1.3.27] | K01095 | AxSpA | 2.92 | 0.01 |
| epsG; transmembrane protein EpsG | K19419 | AxSpA | 2.75 | 0.03 |
| eptC; heptose-I-phosphate ethanolaminephosphotransferase [EC:2.7.8.-] | K19353 | AxSpA | 2.92 | 0.01 |
| nrfA; nitrite reductase (cytochrome c-552) [EC:1.7.2.2] | K03385 | AxSpA | 2.77 | 0.00 |
| ahpF; NADH-dependent peroxiredoxin subunit F [EC:1.8.1.-] | K03387 | AxSpA | 2.98 | 0.05 |
| tarJ; ribitol-5-phosphate 2-dehydrogenase (NADP+) [EC:1.1.1.405] | K05352 | AxSpA | 2.90 | 0.02 |
| K07025; putative hydrolase of the HAD superfamily | K07025 | AxSpA | 3.80 | 0.03 |
| E1.1.1.67, mtlK; mannitol 2-dehydrogenase [EC:1.1.1.67] | K00045 | AxSpA | 2.41 | 0.04 |
| E3.2.1.4; endoglucanase [EC:3.2.1.4] | K01179 | AxSpA | 3.49 | 0.03 |
| arsA, ASNA1, GET3; arsenite/tail-anchored protein-transporting ATPase [EC:7.3.2.7 7.3.-.-] | K01551 | AxSpA | 2.36 | 0.04 |
| tcyL; L-cystine transport system permease protein | K16958 | AxSpA | 2.45 | 0.00 |
| tcyM; L-cystine transport system permease protein | K16959 | AxSpA | 2.40 | 0.01 |
| tcyK; L-cystine transport system substrate-binding protein | K16957 | AxSpA | 2.37 | 0.01 |
| blt; MFS transporter, DHA1 family, multidrug resistance protein | K08153 | AxSpA | 2.41 | 0.04 |
| araP; arabinosaccharide transport system permease protein | K17235 | AxSpA | 2.43 | 0.01 |
| K06921; uncharacterized protein | K06921 | AxSpA | 3.26 | 0.04 |
| K09155; uncharacterized protein | K09155 | AxSpA | 2.73 | 0.04 |
| mexK; multidrug efflux pump | K18303 | AxSpA | 2.36 | 0.01 |
| pabAB; para-aminobenzoate synthetase [EC:2.6.1.85] | K13950 | AxSpA | 2.49 | 0.01 |
| dinD; DNA-damage-inducible protein D | K14623 | AxSpA | 3.07 | 0.01 |
| eta; exfoliative toxin A/B | K11041 | AxSpA | 2.19 | 0.04 |
| vanSB, vanS, vanSD; two-component system, OmpR family, sensor histidine kinase VanS [EC:2.7.13.3] | K18345 | AxSpA | 2.58 | 0.04 |
| MAN; mannan endo-1,4-beta-mannosidase [EC:3.2.1.78] | K19355 | AxSpA | 2.72 | 0.05 |
| tonB; periplasmic protein TonB | K03832 | AxSpA | 3.42 | 0.04 |
| entB; probable enterotoxin B | K11060 | AxSpA | 2.61 | 0.00 |
| tcyN; L-cystine transport system ATP-binding protein [EC:7.4.2.1] | K16960 | AxSpA | 2.38 | 0.01 |
| BLMH, pepC; bleomycin hydrolase [EC:3.4.22.40] | K01372 | AxSpA | 3.17 | 0.02 |
| K10120, msmE; fructooligosaccharide transport system substrate-binding protein | K10120 | AxSpA | 2.49 | 0.00 |
| cyaB; adenylate cyclase, class 2 [EC:4.6.1.1] | K05873 | AxSpA | 2.88 | 0.04 |
| cpdB; 2',3'-cyclic-nucleotide 2'-phosphodiesterase / 3'-nucleotidase [EC:3.1.4.16 3.1.3.6] | K01119 | AxSpA | 3.07 | 0.04 |
| bcrC; undecaprenyl-diphosphatase [EC:3.6.1.27] | K19302 | AxSpA | 3.52 | 0.05 |
| adc; acetoacetate decarboxylase [EC:4.1.1.4] | K01574 | AxSpA | 2.43 | 0.01 |
| proX; glycine betaine/proline transport system substrate-binding protein | K02002 | AxSpA | 2.55 | 0.02 |
| TC.OOP; OmpA-OmpF porin, OOP family | K03286 | AxSpA | 2.86 | 0.04 |
| lctP; lactate permease | K03303 | AxSpA | 2.70 | 0.03 |
| dacB; serine-type D-Ala-D-Ala carboxypeptidase/endopeptidase (penicillin-binding protein 4) [EC:3.4.16.4 3.4.21.-] | K07259 | AxSpA | 3.12 | 0.02 |
| cbiL; nickel transport protein | K16915 | AxSpA | 2.34 | 0.05 |
